# Revealing human sensitivity to a latent temporal structure of changes

**DOI:** 10.1101/2022.06.05.494870

**Authors:** Dimitrije Marković, Andrea M.F. Reiter, Stefan J. Kiebel

**Affiliations:** Department of Psychology, Technische Universität Dresden, 01062 Dresden, Germany; Centre for Tactile Internet with Human-in-the-Loop (CeTI), Technische Universität Dresden, 01062 Dresden, Germany; Department of Child and Adolescence Psychiatry, Psychosomatics and Psychotherapy, Centre of Mental Health, University Hospital Würzburg, Germany; German Center of Prevention Research on Mental Health, Julius-Maximilians Universität Würzburg, Germany

## Abstract

Precisely timed behaviour and accurate time perception plays a critical role in our everyday lives, as our well-being and even survival can depend on well-timed decisions. Although the temporal structure of the world around us is essential for human decision making, we know surprisingly little about how representation of temporal structure of our everyday environment impacts decision making. How does the representation of temporal structure affect our ability to generate well-timed decisions? Here we address this question by using a well-established dynamic probabilistic learning task. Using computational modelling, we found that human subjects’ beliefs about temporal structure are reflected in their choices to either exploit their current knowledge or to explore novel options. The model-based analysis illustrates a large within-group and within-subject heterogeneity. To explain these results, we propose a normative model for how temporal structure is used in decision making, based on the semi-Markov formalism in an active inference framework. We discuss potential key applications of the presented approach to the fields of cognitive phenotyping and computational psychiatry.

## Introduction

The passage of time is a fundamental aspect of human experience. Our behaviour is tightly coupled to our estimate of the elapsed time and the expectations about the time remaining to fulfil short or long-term goals. We are highly sensitive to the temporal structure of our everyday environment and capable of forming precise beliefs about the duration of various events (e.g. a theatre play, traffic lights, waiting in a queue). In practice, temporal structure is typically latent (e.g. not reflected in external clocks) and we seem to rely on an internalised timing mechanism, such as various implicit clocking mechanisms (***Buhusi and Meck, 2005***). This enables us to provide temporal context and an order to events, and to form beliefs about the underlying temporal structure ***Eichenbaum*** (***2014***). It has been proposed that these temporal beliefs are used to make predictions and to adapt our behaviour successfully to ever-changing conditions (***Griffiths and Tenenbaum, 2011***). Therefore, understanding how we learn and represent the temporal structure of our every day environment (***Kiebel et al., 2008***) and use these representations for making decisions (***Marković et al., 2019***) is essential for understanding human adaptive behaviour (***Purcell and Kiani, 2016***).

Neuronal and behavioural mechanisms of time perception have been studied in humans and animals, traditionally using interval timing tasks (***Eagleman, 2008; Meck, 1996***). The key insights of these experiments are that humans and animals integrate the experience of between event duration, in a given context, to form beliefs about possible future duration they might experience. They use these beliefs when estimating or reproducing a newly experienced interval (***Jazayeri and Shadlen, 2010***); in line with a Bayesian account of decision-making (***Shi et al., 2013***). However, it is still an open question how we integrate time perception and beliefs about durations into everyday decision making. Recently, distinct but interlinked research fields have illustrated the importance of temporal representations for cognition and decision making in sequential and dynamic tasks (***Nobre and Van Ede, 2018***; ***McGuire and Kable, 2012***; ***Vilà-Balló et al., 2017***; ***Eichenbaum, 2014***). The sequential neuronal activity in the hippocampus has been suggested to represent elsapsed time (***Eichenbaum, 2017***; ***Buzsáki and Llinás, 2017***; ***Friston and Buzsáki, 2016***), which have led to the postulate of time cells in the hippocampus (***Eichenbaum, 2014***; ***MacDonald et al., 2014***; ***Itskov et al., 2011***) critical for memory and decision-making. For example, in the research on temporal aspects of attention it has been demonstrated that temporal expectations guide allocation of attentional resources in time (***Nobre and Van Ede, 2018***). Similarly, inter-temporal choices or one’s willingness to wait for higher reward is strongly influenced by temporal expectations (***McGuire and Kable, 2012***).

Motivated by the reach literature on temporal representations in the brain, here we focus on the question of how humans form complex temporal representation of their environment. We test how such temporal representations support decisions about whether to explore or to exploit in anticipation of a change in the environment. We introduce a novel computational model of behaviour that describes learning of a latent temporal structure of a dynamic task environment in the context of sequential decision making. The computational model is applicable to any task that can be cast as a dynamic multi-armed bandit problem (***Gupta et al., 2011***) with semi-Markovian changes or switches in the underlying latent states (***Janssen and Limnios, 2013***). Here we specifically apply the model to describe learning in a sequential (probabilistic) reversal learning task (***Costa et al., 2015***; ***Reiter et al., 2016, 2017***; ***Vilà-Balló et al., 2017***). We do so by manipulating temporal contexts in this task: Subjects encountered semi-regular intervals between contingency reversals in one environment. Their behaviour was contrasted with behaviour in another environment where intervals between contingency reversals were irregular.

The proposed behavioural model was based on three components: (i) a set of templates representing possible latent temporal structure of reversals using an implicit representation of between reversal duration (***Yu, 2015***), (ii) the update of beliefs about states and temporal templates derived via approximate inference (***Parr et al., 2019***; ***Yu and Kobayashi, 2003***), and (iii) the action selection, that is the planning process, cast as active inference (***Friston et al., 2017***; ***Markovic et al., 2021***). Together these components allow us to define an efficient and approximate active learning and choice algorithm of latent temporal structures based on variational inference ***Blei et al. (2017)***. Here we extend on our previous investigation of human behaviour in temporally structured dynamic environments (***Marković et al., 2019***). In this work, we demonstrated that a computational model which infers a between-event duration, can be used to reveal subjects’ beliefs about the latent temporal structure in a dynamic learning task. However, a question that has remained open is how humans acquire temporal structure in the first place. Understanding the learning of temporal structure is critical for revealing between-individual variability in temporal expectations and capturing the evolution of temporal representations within individuals. Critically, with the extended model we present here, we are indeed able to capture the learning of temporal representation and address the non-stationarity of subjects’ temporal representation during the course of the experiment.

Our aim is to address the following questions: (i) Are subjects a priori biased towards expecting regular or irregular temporal structure? (ii) Are subjects able to learn latent temporal structure without explicit instructions? (iii) How does the quality of temporal representation impact their performance? Using simulations we can illustrate the interaction of accurate representation of temporal structure and behaviour, mainly performance on the task and the engagement with exploratory behaviour. Using model-based analysis we demonstrate high diversity between subjects both in their prior beliefs about temporal structure, and their ability to adapt their beliefs to different latent temporal structure. Critically, we link the quality of temporal representation to subjects’ performance both in terms of group-level performance and within-subject variability of their performance during the task.

In what follows we will first briefly describe the experimental task, provide the overall summary of behavioural characteristics, introduce the behavioural model, and finally show results of the model-based analysis of behaviour. The formal details of the approach are described in Methods and Materials.

## Results

A typical probabilistic reversal learning task asks subjects to make a binary choice between two options, e.g. A and B, where each option is associated with a probability of receiving a reward or punishment. For example, initially choosing A returns a reward with a high probability *p_H_* = 0.8 and choosing B returns a reward with low probability *p_L_* = 0.2. Importantly, after several trials the reward contingencies reverse, i.e. switch, such that choosing B returns the reward with high probability *p_H_*. However, subjects are not informed about the reversal and they have to infer that a change occurred from the feedback they receive in order to adapt their behaviour. From the point of view of participants, a reversal can be difficult to detect as outcomes are probabilistic. This means that if someone observes a loss after a sequence of gains, e.g. when choosing the option A, this could be caused either by: (i) a true reversal, where now option B is rewarded with the probability *p_H_* or (ii) by an unlucky outcome of an otherwise correct choice. To obtain a more direct information about the subjective uncertainty of participants about the correct choice (i.e. choosing the option with high reward probability, *p_H_*) on any given trial, we extended the standard design with an additional third exploratory option. This new option does not result in monetary gain or loss but provides information about the correct choice on a current trial. A high uncertainty about the best choice (current context) can be easily resolved by selecting the epistemic option. We will label all choices of the exploratory options as exploratory, and all other choices as exploitative (note that the outcomes of exploitative options also provide some information about the current context). A trial sequence of the experimental task is shown in Figure 1.

**Figure 1.**
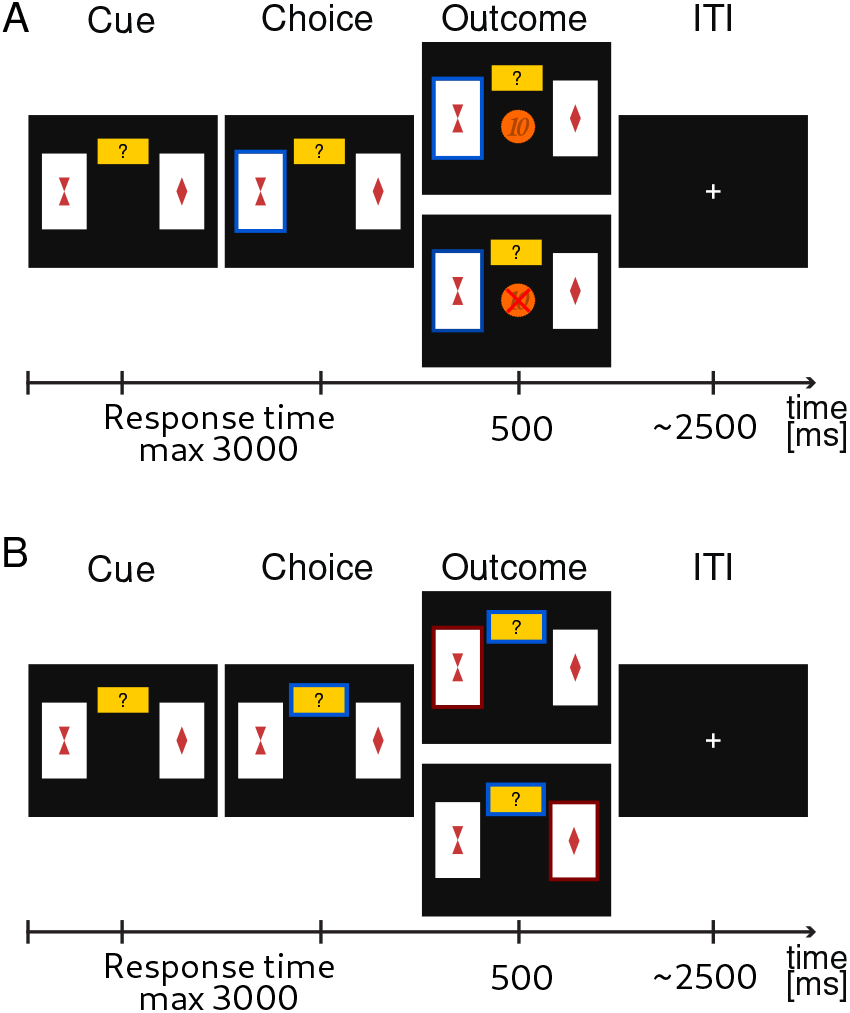
Probabilistic reversal learning task. Exemplary trial sequence. Subjects were instructed that one card always had a higher probability of a monetary reward. They were instructed to choose the card that they thought would lead to a monetary gain with higher probability, or, alternatively, choose to explore (small yellow rectangle with question mark). The latter would provide them with a correct information about what option would have had a higher probability of reward **A**. If participants had chosen one of the cards, the corresponding card was highlighted and feedback was displayed. The feedback consisted of either the visual display of a 10 Euro cents coin in the centre of the screen for a gain outcome, or a crossed 10 Euro cents for a loss outcome. **B**. If the participant had chosen the explore option, the card with the currently highest reward probability was highlighted (either the left or the right card).

To investigate subjects’ ability to learn latent temporal structure we defined two ex-perimental conditions (manipulated in a between-subject design), one with irregular reversals and another with regular reversals (see Figure 2). In the condition with irregular reversals, the moments of reversals are not predictable and between-reversal intervals are drawn from a geometric distribution (Figure 2A). In the condition with regular reversals, the moments of reversal are predictable, and they occur at semi-regular intervals, drawn from a negative binomial distribution (Figure 2B). Subject were randomly assigned to one of the two possible conditions, as illustrated in Figure 2. In the first condition, subjects experience irregular reversal statistics for 800 trials, after which the reversals occur at semi regular intervals for the last 200. In the second condition, subjects experience semi-regular reversal statistics for 800 trials, and then the irregular reversal statistics during the last 200 trials. Note that when changing the temporal statistics we copied the time series of reversals from the initial 200 trials of the different condition.

**Figure 2.**
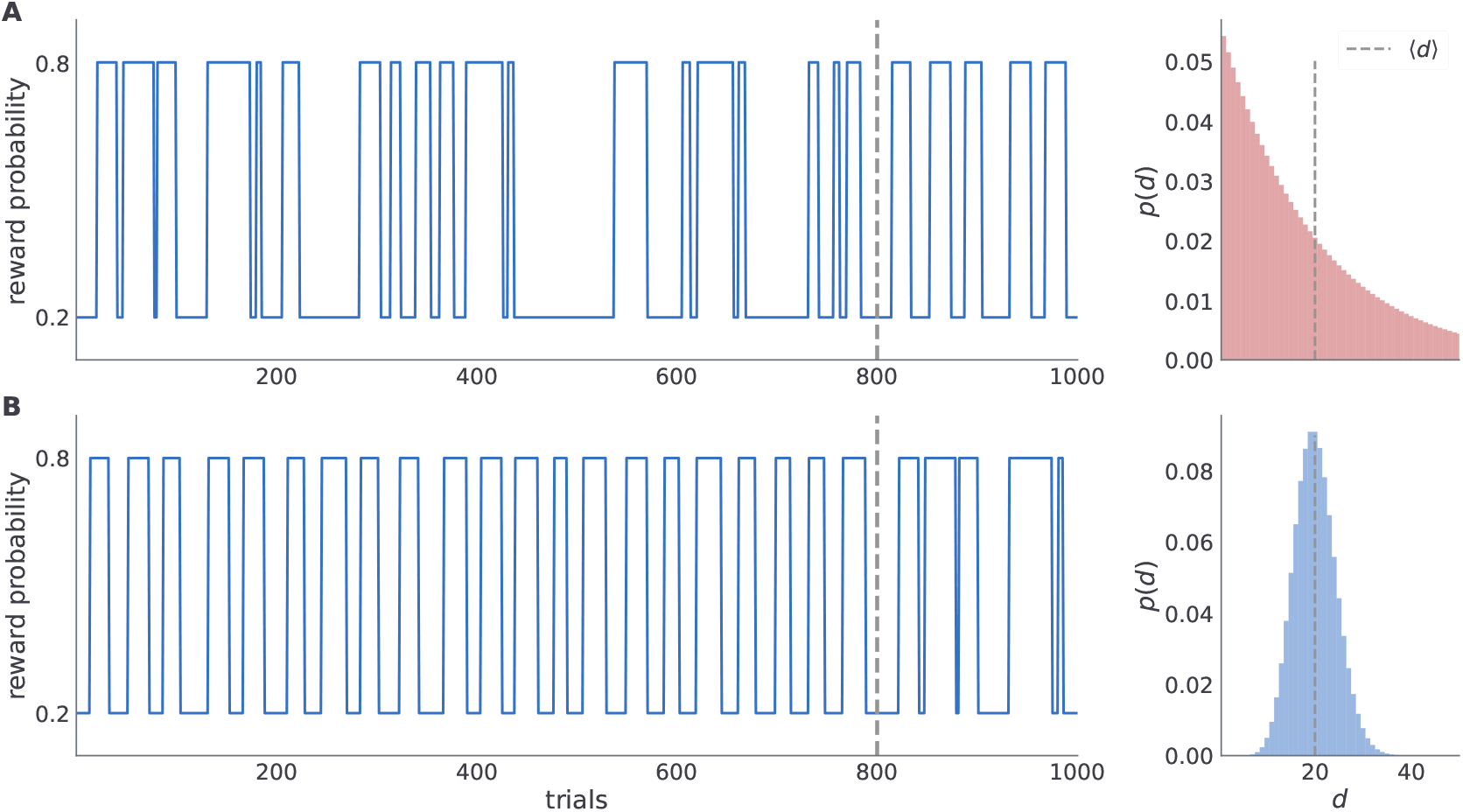
Time series of reward probabilities. A, Condition with irregular reversals, and B, condition with (semi-)regular reversals. The reward probability of the high-probability stimuli at any time step was set to *p_H_* = 0.8 and the low-probability stimuli to *p_L_* = 0.2. Dashed vertical lines shows the moment of change of the latent temporal structure: (i) from irregular to semi-regular statistics in the irregular condition, and (ii) from the semi-regular to irregular statistics in the regular condition. Figures on the right illustrate the generative distribution of the between-reversal intervals *d* for each condition. Note that the mean between reversal duration 〈*d*〉 is identical in both conditions.

Importantly, the moments of reversal were fixed within each group (depending on the condition reversals occur always on the same trials), and the same choice by different subjects on any given trial would lead to the same outcome (the outcome statistics were generated only once for each condition and trial, and then replayed to all subjects depending on their choices and the condition they were assigned to).

### Analysis of choice data

We will first describe the behavioural characteristics of the two groups of subjects exposed to the two different experimental conditions. The two behavioural measures of interests here are the *performance* (odds of being correct, i.e. odds of choosing the option with the higher reward probability) and *probing* (odds of exploring, i.e. odds of choosing the exploratory option). We describe all the behavioural measures in detail in Behavioural measures subsection of Methods and Materials.

Subjects (*N* = 74) were pseudo-randomly assigned to one of the two experimental conditions, where *n_r_* = 41 participants were assigned to the condition with regular reversals, and *n_i_* = 33 to the condition with irregular reversals. Note that some subjects rarely engaged with exploratory option. Out of 50 subjects who where exposed to the variant of the experiment with exploratory option (24 subjects performed a standard version of the task without exploratory option, see Experiment for more details), 5 subjects never engaged with the exploratory option. In Figure 3 we provide a summary of average behavioural measures for individual subjects. We do not find any significant performance differences between the two regularity conditions, see Figure 3A. However, for the subset of subjects which interacted with the exploratory option (45 subjects) we find that the performance is positively correlated with probing (Pearson correlation coefficient *r* = 0.6, with *p* < 10^-4^), see Figure 3B. Interestingly, neither of the two behavioural measures (when plotted as a within subject average over the course of experiment), reveals obvious between-condition differences. However, when comparing the temporal profile of these measures over the course of experiment (see ***Figure 9-Figure Supplement 1***), one notices large variability both between subjects but also within a subject over the course of experiment;suggesting ongoing learning of the task structure. In what follows we will classify the heterogeneity of behavioural responses using a model-based analysis.

**Figure 3.**
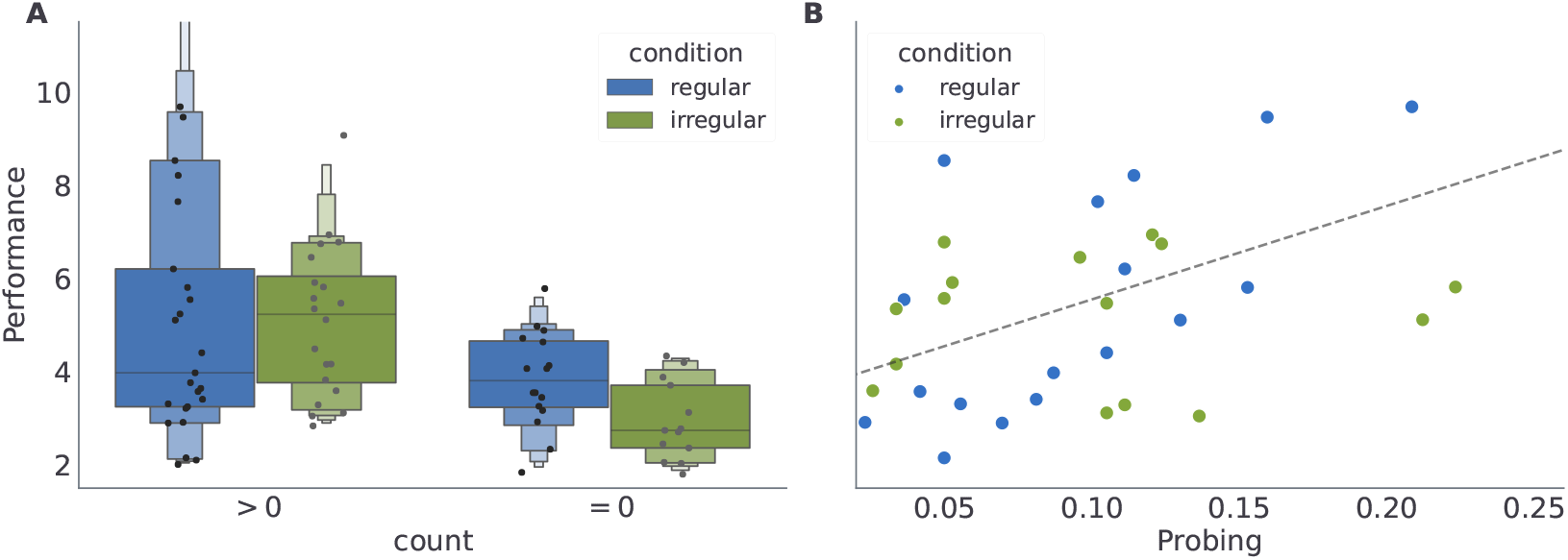
Averages of behavioural summary measures. A, Distribution of the mean performance of subjects with low and high number of exploratory choices, see methods. B, Dependence of the mean performance on the mean probing, where we excluded participants without exploratory choices (count = 0). Note that when computing mean performance and mean probing for each participants, we have excluded the first 400 (initial responses during which the subjects might have still been adjusting to the task) and last 200 (responses after the change in the reversal statistics) responses of each participant, see Methods and Materials - Model inversion for the motivation for the cutoff.

### Behavioural model

The behavioural model will allow us to investigate the process of learning of the latent temporal structure in different experimental conditions, reveal subjects’ preferences to engage with exploratory option (collect information), and subjects’ motivation to collect rewards. We achieve this by fitting free model parameters to behavioural responses of each subject (see Methods and Materials for more details). Our aim with the model based analysis is to quantify beliefs about temporal structure of reversals and understand how the belief dynamics influences subjects’ behaviour.

We conceptualised the behavioural model as an active inference agent (***Friston et al., 2015, 2016***) with hidden semi-Markov models (***Yu, 2010***), which is capable of representing and inferring latent temporal structure. In active inference, besides defining perception and learning as a Bayesian inference process, action selection is also cast as an inference problem aimed at minimising the expected surprise about future outcomes, that is, the expected free energy (***Smith et al., 2022***) (see also Eq (18)). Through its dependence on the expected free energy, the action selection has an implicit dual imperative (see possible factorisation of the expected free energy in Eq (18)): The expected free energy combines intrinsic and extrinsic value of a choice, where intrinsic value corresponds to the expected information gain, and the extrinsic value to the expected reward of different choices. The implicit information gain or uncertainty reduction pertains to beliefs about the task’s dynamical structure and choice-outcome mappings, e.g. (***Kaplan and Friston, 2018***; ***Schwartenbeck et al., 2013***). Therefore, selecting actions that minimise the expected free energy dissolves the exploration-exploitation trade-off, as every action is driven both by expected value and the expected information gain. This is a critical feature of active inference models which allows us to account for exploratory choices (see Figure 1).

We express the agent’s generative model of task dynamics in terms of hidden semi-Markov models (HSMM) (***Yu, 2015; Marković et al., 2019***). The HSMM framework extends a standard hidden Markov model with an implicit (or explicit) representation of durations between consecutive state changes. HSMM have found numerous applications in the analysis of non-stationary time series in machine learning (***Duong et al., 2005; Gales and Young, 2008***), and in neuroimaging (***Borst and Anderson, 2015***; ***Shappell et al., 2019***). HSMM have also been used in decision making for temporal structuring of behavioural policies (***Bradtke and Duff, 1994***) or in temporal difference learning as a model of dopamine activity when the timing between action and reward is varied between experimental trials (***Daw et al., 2002***).

Here, we use the semi-Markov representation of task dynamics within the behavioural models to define an agent that can learn latent temporal structure, form beliefs about moments of change, and anticipate state changes. We implemented the learning of the hidden temporal structure of reversals as a variational inference scheme, where we assume that the agent entertains a hierarchical representation of the reversal learning task, with a finite set of models of possible temporal structure of the dynamic environment. In other words, we assume that human brain entertains a set (possibly a very large set) of temporal templates. In Figure 4, we show the graphical representation of the generative model of behaviour, which is described in detail in Behavioural model section. Here we will briefly introduce the relevant parametrisation of the behavioural model, which are critical for understanding the model comparison results presented in the next subsection.

**Figure 4.**
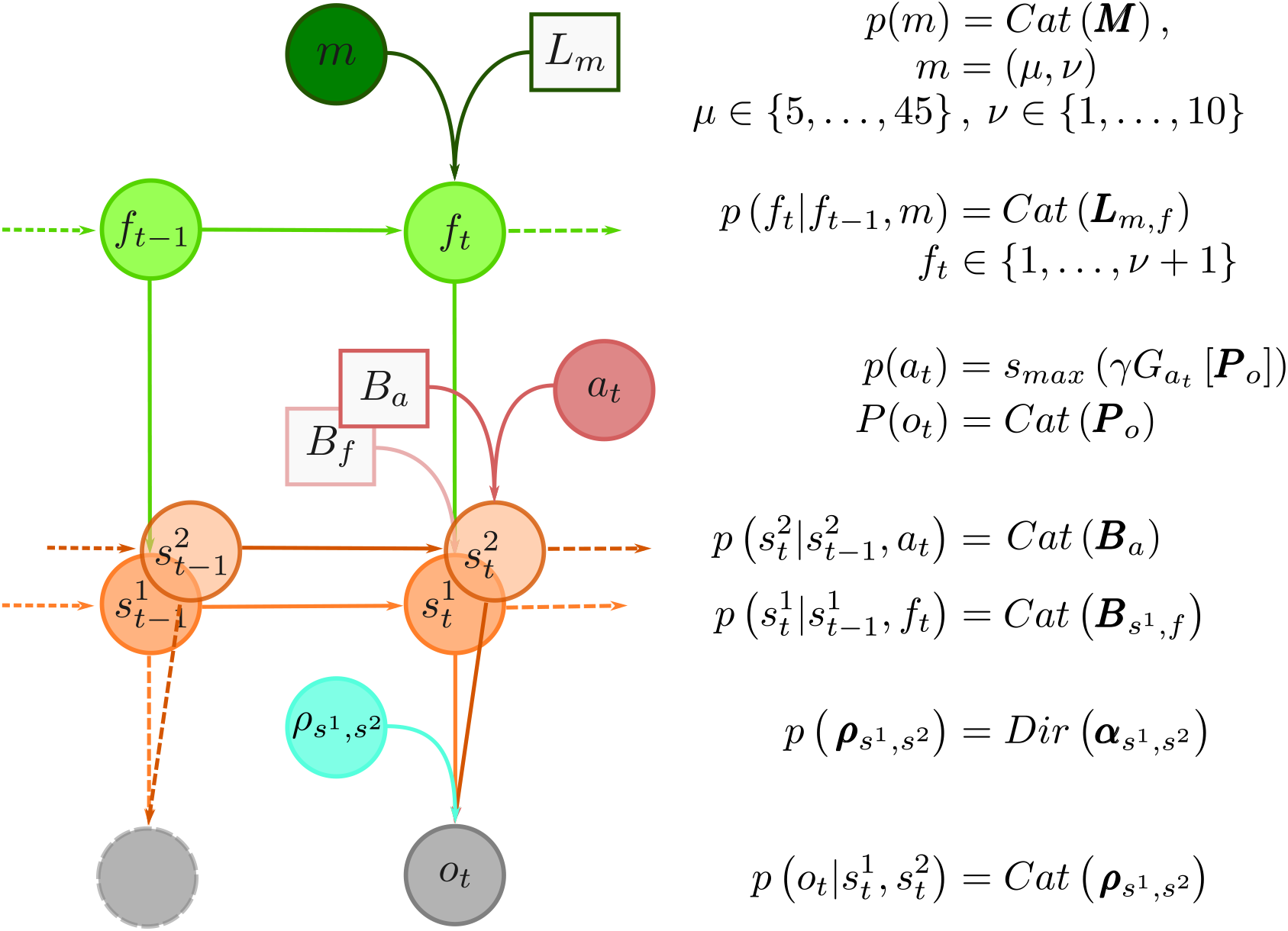
Graphical representation of the generative model and model summary. At the top of the hierarchy is the temporal template variable *m*. The total number of temporal templates is finite, e.g. *m* ∈ {1,…, *m_max_*}, and each template *m* provides an implicit representation of a prior probability distribution over between-reversal intervals *d*, parameterised with a pair *m* = (*μ, v*), where *μ* expresses mean between reversal interval, and *v* plays a role of a precision parameter, that defines regularity of between-reversal intervals. The implicit representation of temporal structure is encoded with probability transition matrices ***L**_m_* latent phases *f*. The number of latent phases depends on precision *v*. The reversal can occur only when end phase is reached (*f* = *v* + 1). Therefore, the phase variable *f* controls the transitions probability ***B***_*s*^1^,*f*_ between latent states (the correct choice) of the task denoted as random variable *s*^2^ ∈ {1, 2}. At every trial *t* the subject makes a choice *a_t_* hence decides on an option *s*^1^ which will generate an outcome *o_t_*. The choices are deterministic, hence 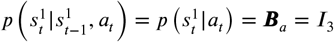. Finally, the choice-outcome contingencies are treated as latent variables ***R***_*s*^1^,*s*^2^_ which have to be learned over the course of the experiment. We use a vague Dirichlet prior over choice-outcome contingencies. Inverting the generative model of outcomes using variational inference defines the inference and learning component of the behavioural model. In turn, marginal beliefs about latent states 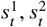 and parameters ***R***_*s*^1^, *s*^2^_ are used to define action selection, that is compute the choice likelihoods using the expected free energy (Eq (18)).

Each temporal template *m* corresponds to a pair of parameters *m* = (*μ, v*) that define the frequency of reversals *μ* and the regularity of reversals *v* (the higher the value the more regular the changes are). In Figure 12 we illustrate three of these templates, which differ in their regularity parameter *v*, but all have the same frequency parameter *μ*. It is important to note that when *v* = 1 (the lowest value) the temporal templates correspond to the hidden Markov model (HMM) representation. HMM representation implies that the moments of reversals are unpredictable, or maximally irregular. Here we use the HMM representation as a reference point for determining whether participants were able to learn latent temporal structure of reversals, and whether they a priori expected predictable moments of reversal.

When simulating behaviour and fitting the model to participants’ choices, we use a prior probability *p* (*m*) over temporal templates *m* to restrict otherwise rich set of all possible temporal templates *m* = (*μ, v*), that span all combinations of *μ* ∈ {5,…, 45} and *v* ∈ {1,…, 10}. Hence, template prior *p* (*m*) reflects prior expectations of an agent at the beginning of the experiment about the possible temporal structure of the task dynamics. Therefore, to capture a wide range of prior beliefs we require a flexible prior *p*(*m*) that can reflect subjects with different prior expectations about temporal structure. Posterior estimates of the most likely parametrisations of the temporal prior, allows us to infer from the behavioural data if participants’ beliefs are a priori precise and biased towards expecting irregular reversals, or are imprecise and accommodate a wide range of possible latent temporal structures. In the model, we use the following prior over temporal templates:

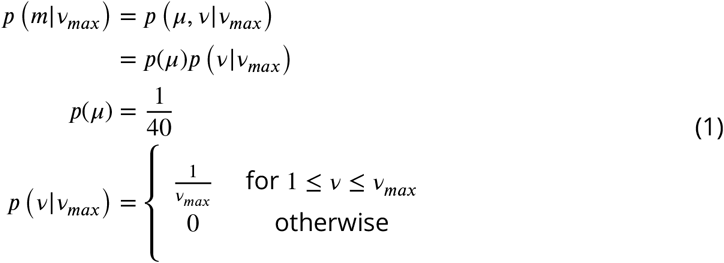

where *v_max_* ∈ {1,…, 10}. Note that the prior regularity parameter *v_max_* reflects Bayesian prior expectations about the maximal precision of between-reversal intervals. In other words, *v_max_* captures the agent’s expectations about the maximal regularity of reversals, and hence their predictability. Thus, with this parametrisation we assume that subjects, at the beginning of the experiment, have uniform beliefs about a possible mean duration between reversal interval, but might differ in their propensity to represent high or low regularity of between-reversal intervals. For example, some subjects could hold precise beliefs that reversals were not under their control and were therefore inherently unpredictable (corresponding to *v_max_* = 1). Such a subject would fail to learn - or accumulate evidence for - the regularity of reversals in the regular condition. Conversely, some participants may have imprecise prior beliefs about regularity (***v_max_*** > 1);enabling them to learn that reversals were regular, thus predictable, in the appropriate condition.

The beliefs about temporal templates, influence the beliefs about the reversal probability on any given trial (i.e., how likely is that a reversal occurs in the next trial), and consequently modulate beliefs about the latent state of the task (i.e., which card is associated with high reward probability) and corresponding outcome probabilities. In turn, the beliefs about the latent state influence the choices. As mentioned above, choices are defined as the minimisers of the expected free energy (surprise about future outcomes), typically denoted by *G*. Given the expected free energy *G_a_* [***P**_o_, v_max_, t*] of action *a* on trial *t* we define the choice likelihood as

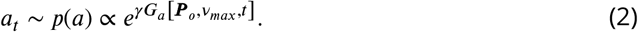

Here, the parameter *γ* denotes choice precision, thevectorof probabilities 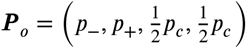 denotes prior preferences over possible outcomes, that is, losses (-), gains (+), and cues (c). In active inference (***Friston et al., 2017***) prior preference parameter ***P**_o_* defines motivation to collect rewards (generate correct choices) and collect information (engage with the exploratory option). In more familiar terminology of reinforcement learning, the logarithm of prior preferences ln ***P**_0_* assigns a subjective value to possible outcomes, and the expectation of log-preferences defines the expected value of different actions (see Eq (18)). Importantly, Eq (2) acts in two different ways: (i) as a mapping from beliefs into actions which we used to simulate behavioural choices, and (ii) as a choice likelihood which we use for inverting the model when fitting the model to subjects’ choices to derive the posterior estimates of free model parameters (*γ*, *p*_−_,*p*_+_, *v_max_*), individually for each subject. Details of the model inversion procedure are described in Model inversion.

#### Simulating the behavioural effect of prior expectations over temporal templates

By simulating the model’s behaviour given different values of temporal regularity parameter *v_max_*, we aimed to demonstrate that the agent can acquire a correct representation of the latent temporal structure in different experimental conditions, and that *v_max_* influences the dynamics of both performance and probing. Importantly, different values of *v_max_* should lead to sufficiently distinct behaviour, if we hope to accurately associate subjects’ behaviour with underlying model parametrisation.

The temporal regularity parameter *v_max_* is the key parameter in the model to understand how learning about temporal structure comes about. As *v_max_* constrains the maximal temporal regularity the agent expects in the task, it is a measure of subjects’ sensitivity to the latent temporal structure. Importantly, we find that varying *v_max_* results in simulated behaviour with distinct behavioural patterns in our two experimental conditions as shown in ***Figure 5***. As we increase *v_max_* the behavioural performance increases, in both conditions. In contrast, as we increase *v_max_* the probing decreases, as the agent is more certain about the moment of reversal and requires information provided by exploratory option less often. Note that different values of *v_max_* induce stronger differences in both performance and probing in the regular condition, compared to the irregular condition. Practically, this means that we can infer *v_max_* from behavioural data with higher precision in regular than in irregular condition. We validate the classification accuracy of *v_max_* based on posterior estimates given simulated data in the form of confusion matrix as shown in ***Figure 8-Figure Supplement 1***. Note that even in the ideal case when behaviour is generated exactly from the behavioural model, classification accuracy with regard to *v_max_* is substantially lower in irregular compared to irregular condition. We will clarify the impact of this in the next subsection when discussing the the results of model-based analysis.

**Figure 5.**
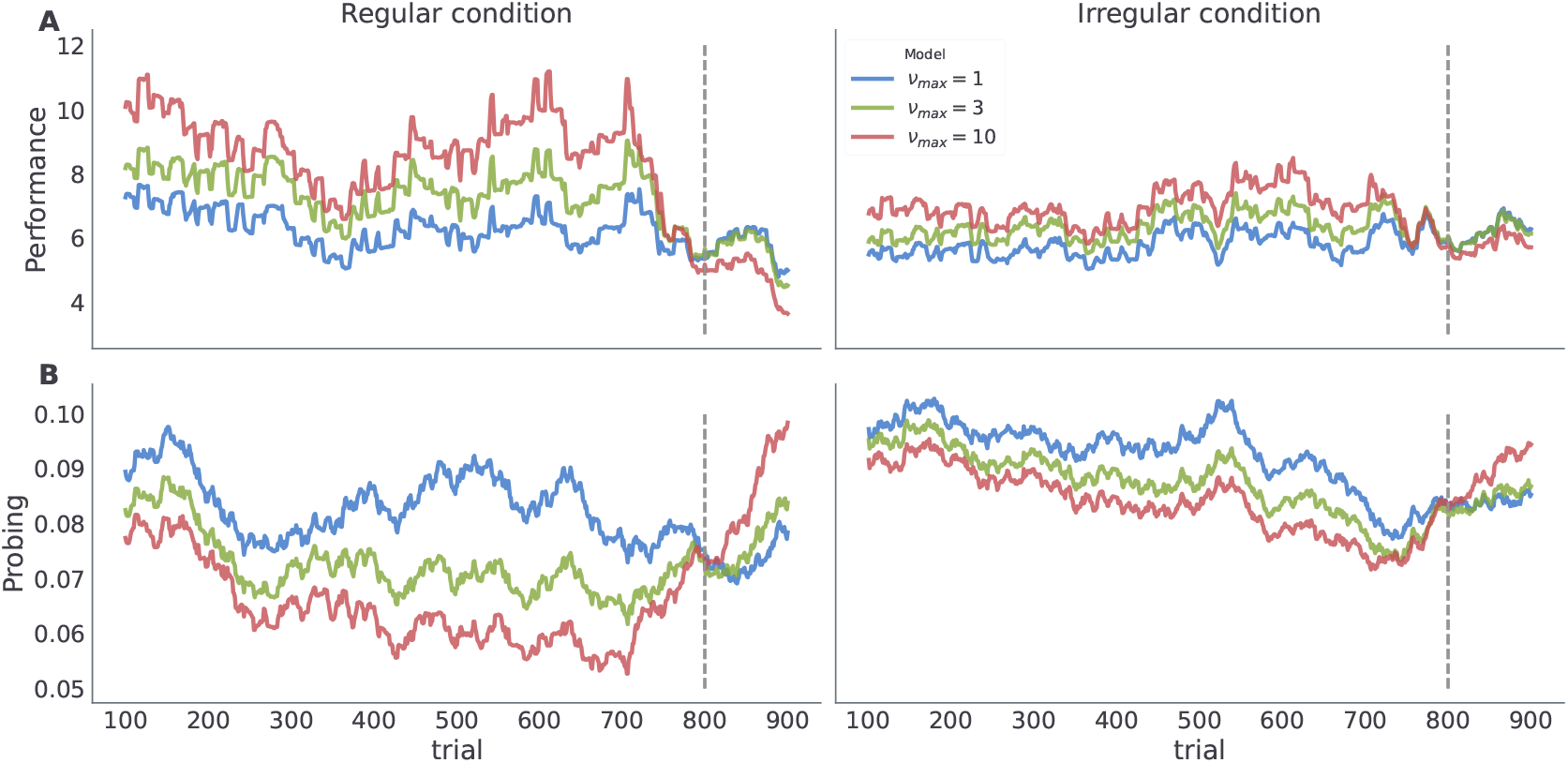
Model dependent dynamics of behavioural measures for varying *v_max_*. Each line corresponds to an average over *n* = 50 simulated trajectories with *γ* = 5, and ***P**_o_* = (0.1, 0.6, 0.15, 0.15). **A**, Performance estimated as odds of generating a correct choice withing a 200 trials long time window centred at trial index. **B**, probing, computed as odds of selecting the exploratory option within the 200 trials long time window.

#### Demonstrating the learnability of latent temporal structure

As a next step we will illustrate that the agent with the highest value of temporal prior (*v_max_* = 10)—that is, the agent with the most adaptable beliefs about the latent temporal structure—is capable of accurately inferring the correct temporal template *m*, and that the rate at which agent learns correct representation of the temporal structure depends on the given temporal context. Hence we expect that human subjects should also be capable of learning the correct statistics with comparable prior expectations about temporal structure. In ***Figure 6*** we show posterior beliefs over temporal templates in the form of marginal posterior beliefs about the mean *μ* and the regularity *v* at each time step of the experiment. We see that the agent quickly learns the correct mean between-reversal duration (already after 200 trials the highest posterior probability is close to *μ* = 19), but it takes longer (more than 400 trials) to form precise beliefs about the level of temporal regularity. In contrast, in the irregular condition, learning the correct mean between-reversal-interval (fixed to *μ* =19 in both conditions) takes more time and is less precise, but the posterior estimates over the precision parameter (*v*) converge faster to the correct values (already after 200 trials). Note that having the correct representation of both mean and precision parameters is more important in the regular condition as one can achieve higher improvements in the performance compared to the irregular condition, as we demonstrated previously in ***Marković et al.*** (***2019***).

**Figure 6.**
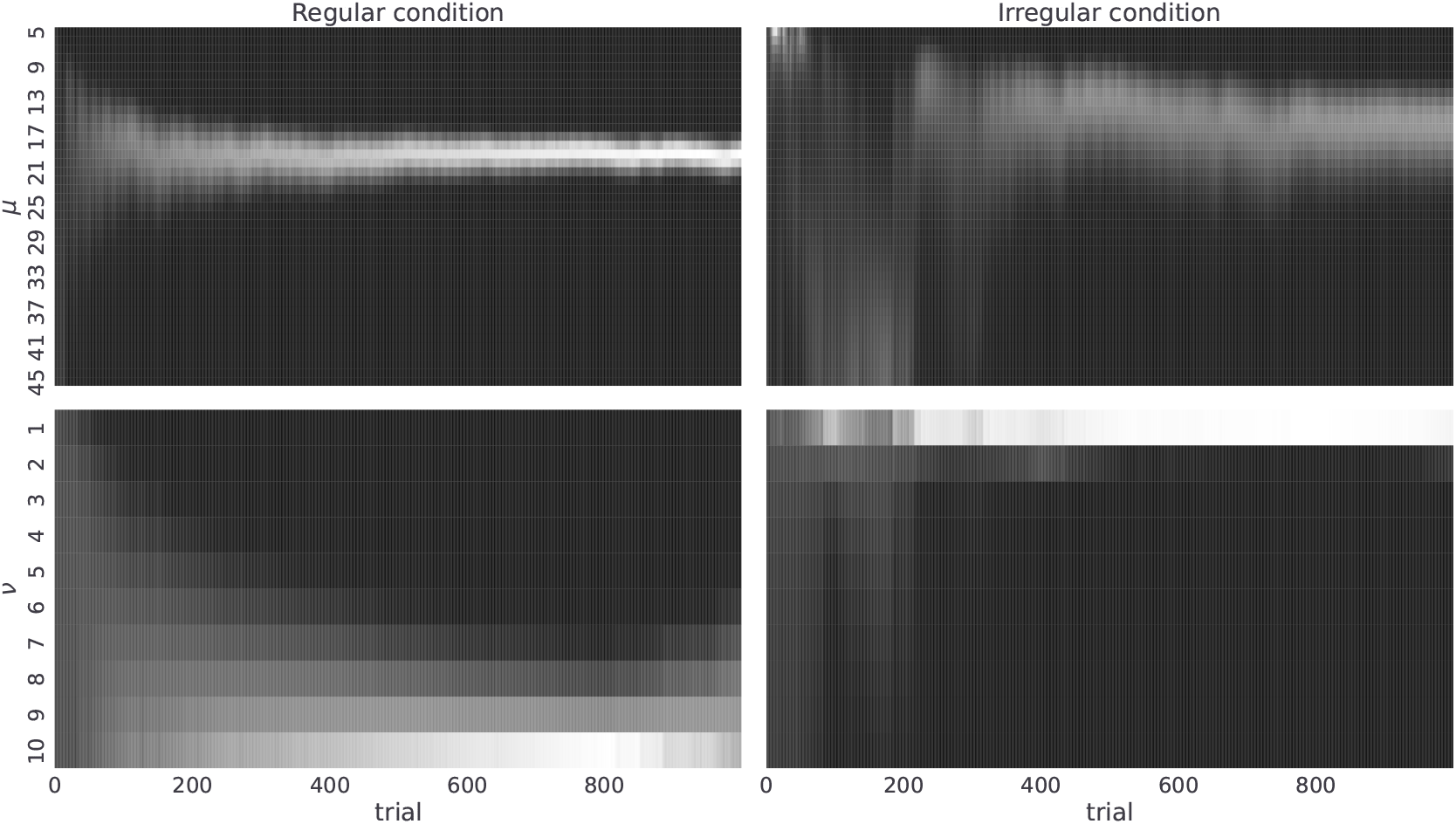
Posterior beliefs about temporal templates. Posterior beliefs of a single agent in the regular (left column) and the irregular condition (right column). Posterior beliefs *q_t_*(*m*) = *q_t_*(*μ, v*) at each trial *t* over templates *m* are marginalised over precision parameter obtaining *q_t_*(*m*) (top) and mean parameter *μ* obtaining *q_t_*(*v*) (bottom). The posterior beliefs are estimates obtained from a single run of the agent in both experimental conditions where we fixed the temporal prior parameter to *v_max_* = 10, choice precision to *γ* = 5, and the preference vector to ***P**_o_* = (0.1, 0.6, 0.15, 0.15). The lighter the colour the higher is the corresponding posterior probability for that parameter value.

#### Simulating the behavioural effect of prior preferences over outcomes

As mentioned above, the prior preference over outcomes ***P**_o_* parameterise agents’ motivation to collect rewards (generate correct choices) and collect information (engage with the exploratory option). Therefore, it is important to understand how prior preferences interact with performance and probing. We show that the more an agent engages with the exploratory options, (i.e., the higher its preference for choice cues), the better its representation of latent temporal structure, and consequently the higher agent’s performance. This is because selecting exploratory options maximally reduces the uncertainty about the latent state (which option has higher reward probability) which in turn allows an agent to learn a more accurate representation of the latent task dynamics. We visualise these dependencies in ***Figure 7***, where we show what impact changing *p*_+_ and *p*_−_ have on performance, probing, and the quality of temporal representation after 800 trials. In the supplementary figure ***Figure 7-Figure Supplement 1*** we show same dependencies but with respect to changing *p*_+_ and *p_c_*, hopefully helping the reader to build an intuition about interactions between prior preference parameter and behaviour. Note that in both figures we only consider cases in which *p*_+_ ≥ *p*_−_ as this reflects higher prior preference for gains than for losses in the agent, which we would expect to hold for all subjects.

**Figure 7.**
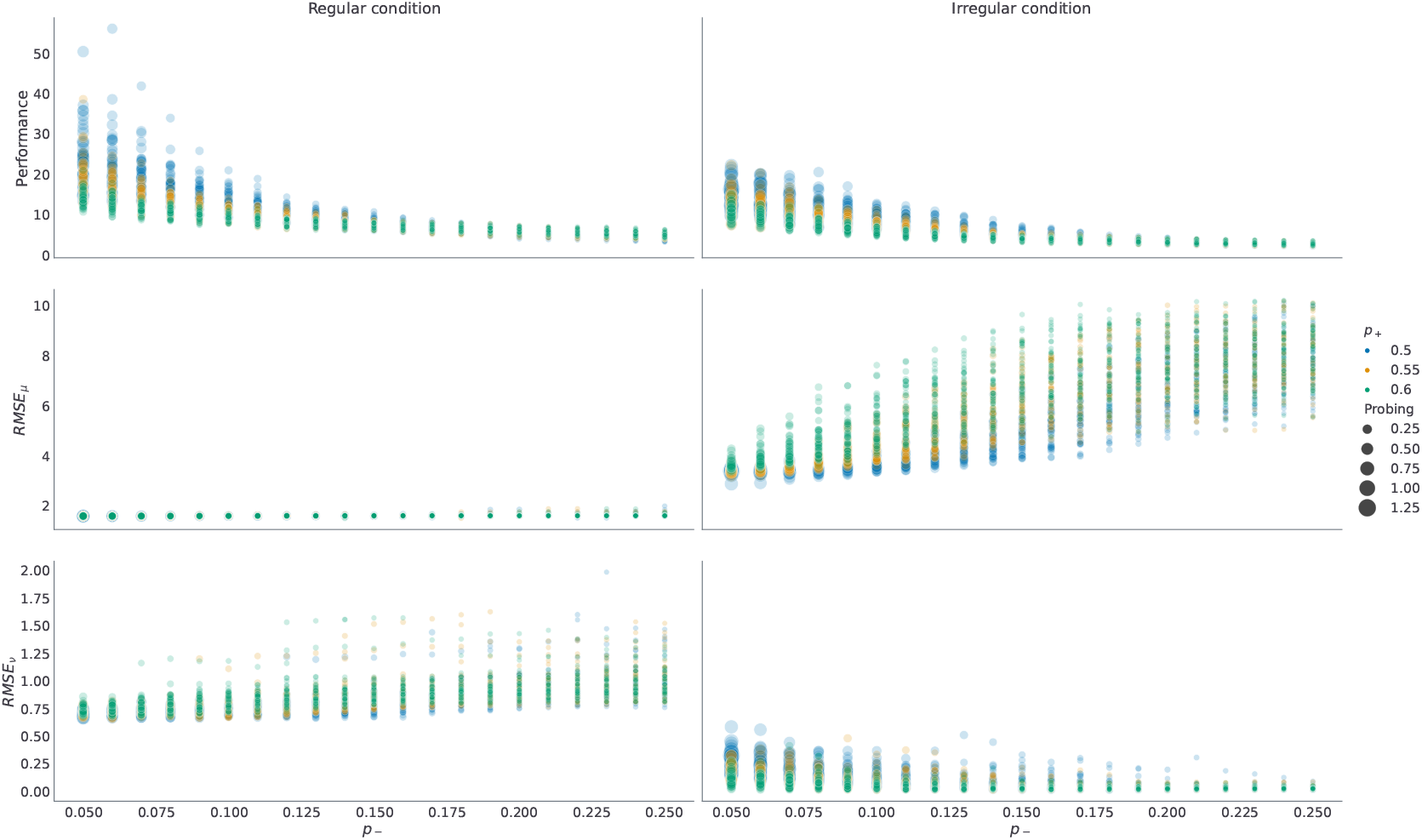
Dependence of performance, probing, and the quality of temporal representations on prior preferences over outcomes. Each dot in the plot corresponds to a single run with fixed prior preferences 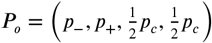, with temporal prior set to *v_max_* = 10, and with decision precision set to *γ* = 5. For each possible pair ((*p*_−_, *p*_+_) ⊗ {0.05, 0.06,…, 0.25} ⊗ {0.5, 0.55, 0.6}) we have repeated *n* = 50 simulations in each condition. Here 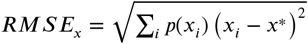 stands for the root mean square error (RMSE) of corresponding parameters *μ, v* that define temporal template. The RMSE is computed using posterior probabilities *q_t_*(*μ, v*) obtained at trial *t* = 800. The performance and the probing are computed as averages over responses from trial *t* = 400 until trial *t* = 800. Note that probing is increasing as we reduce *p*_−_ and keep *p*_+_ fixed (circles of the same colour), and as we reduce *p*_+_ and keep *p*_ fixed, as the larger the sum (*p*_+_ + *p*_) is, the lower is the preference for choice cues *p_c_*, and hence the tendency of the agent to engage with the epistemic option. Higher probing (larger circle size) results in higher performance in both conditions (top row). Similarly, both *RMSE_μ_* and *RMSE_v_* are lower for larger exploration odds, with the exception of *RMSE_v_* in irregular condition. Note that forming an accurate temporal representation is especially important in the regular condition, where forming correct anticipatory beliefs can substantially improve behavioural adaptation and simplify the problem of balancing between exploratory and exploitative choices. In contrast, in the irregular condition, having a precise representation of temporal structure does not impact performance substantially, and the agent performs better when engaging with the epistemic option more often. **Figure 7–Figure supplement 1.** Dependence of performance, probing, and the quality of temporal representations on prior preference over cues *p_c_*.

### Model-based analysis of subjects’choices

Using the model-based analysis of the choice data we next ask whether human subjects can learn latent temporal regularities in the reversal learning tasks? An individual’s capacity to learn correct temporal regularity corresponds to their behaviour being associated with a less precise prior over temporal templates (Eq (1)), that is, larger *v_max_*. An agent with imprecise prior over temporal templates is able to learn an accurate representation of a distribution of between-reversal-intervals, and to form expectations about the moment of reversals (see ***Figure 6*** and ***Figure 7***) in both conditions. Thus, we anticipated that between-subject variability in performance and probing would be reflected in different posterior estimates of the most likely *v_max_* value associated with the behaviour of individual subjects.

Therefore, we first classify subjects based on the maximum a-posteriori estimate over possible values of *v_max_* ∈ {1,…, 10}, as shown in ***Figure 8***. For each subject we compute a posterior probability over *v_max_* and assign the subject the value of the temporal prior *v_max_* corresponding to the value with the highest exceedance probability (see Model inversion). Using this procedure we find that 8 out of 41 subjects in the regular condition, and 17 out of 33 subjects in the irregular condition are assigned to the group with temporal prior *v_max_* > 1. For the subjects in the regular condition this result suggests that they have learned to anticipate reversals to certain extent. However, it is critical to note that the classification of subjects is not perfect, specially in irregular condition as shown in ***Figure 8-Figure Supplement 1***. However, as our aim is not to identify precisely participants’ temporal prior, but simply to distinguish between subjects that learn temporal regularities (*v_max_* > 1) from those that do not (*v_max_* = 1), limiting the analysis to binary classification leads to the following classification accuracy in simulated data: (i) in the regular condition *v_max_* = 1, *ACC* = 1., and *v_max_* > 1, *ACC* = 0.91, (ii) in the irregular condition *v_max_* = 1, *ACC* = 0.4 and *v_max_* > 1, *ACC* = 0.8. Low classification accuracy in the irregular condition is the reason for substantially higher number of subjects assigned to the *v_max_* > 1 model class.

The posterior estimates of model parameters shown in ***Figure 8*** show that 65% of the participants were assigned to the model class corresponding to the simplest HMM representation (*v_max_* = 1) which assumes maximal irregularity. Importantly, when we plot the time course of both performance and probing, as shown in ***Figure 9***, we find the similar trajectory of behavioural measures over the course of the experiment as what we see in simulated data. Namely, that the performance is higher and the probing is lower in the group of participants associated with larger *v_max_* (compare with ***Figure 5*** and ***Figure 5-??***). The result is notably more robust for the regular condition than the irregular. The behavioural trajectories of individual participants are shown in ***Figure 9-Figure Supplement 1***.

**Figure 8.**
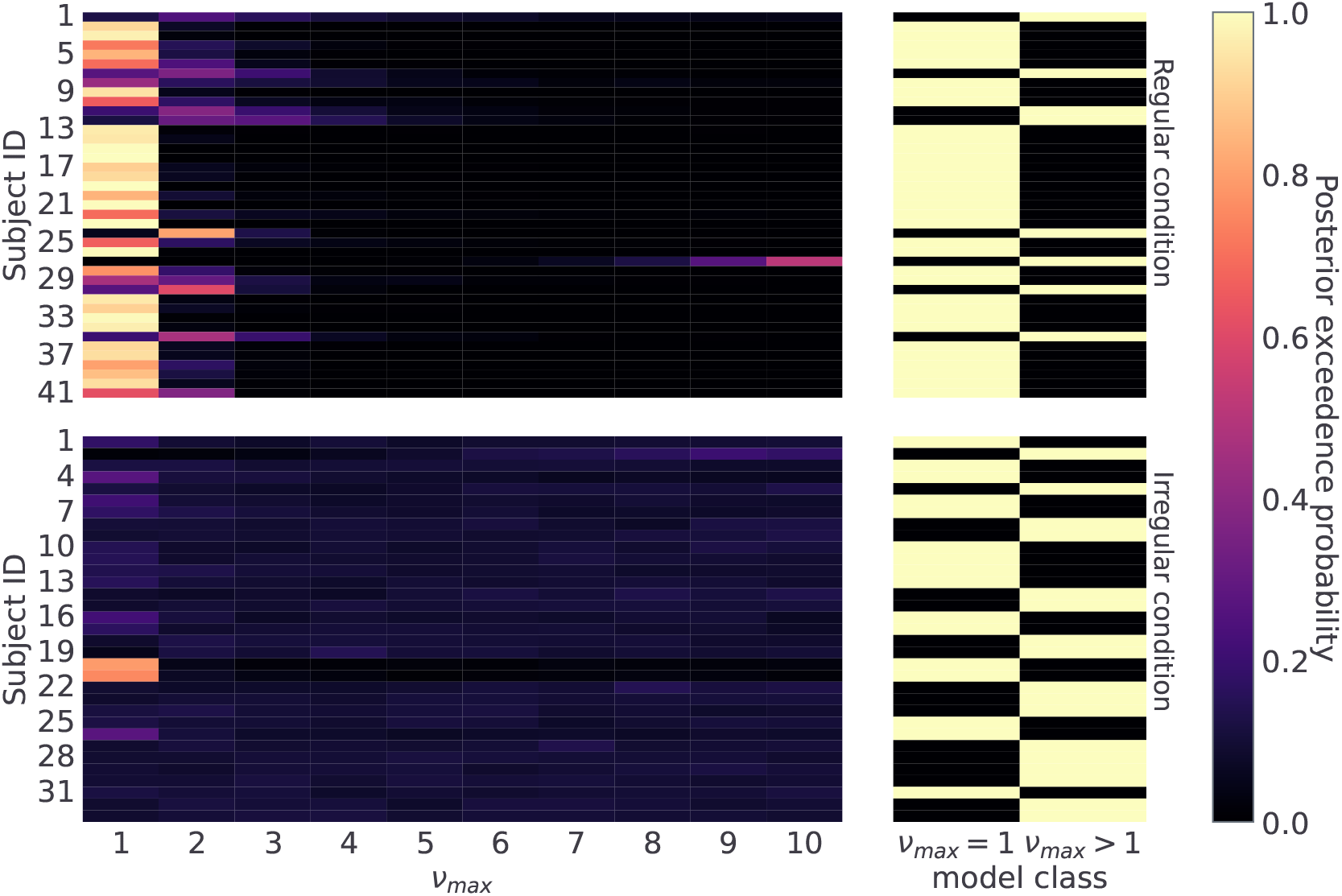
Posterior probability over temporal prior *v_max_*. Posterior probability of possible *v_max_* values for each subject, reflecting a subject’s flexibility to learn latent temporal structure. On the right hand side, we combine posterior estimates into two classes, one for the limiting case *v_max_* = 1, and another for all other options *v_max_* > 1. This split differentiates subjects not sensitive to temporal regularities from the ones who a priori expected a regular temporal structure of reversals. Note that lighter colours correspond to higher posterior probability. **Figure 8–Figure supplement 1.** Confusion matrix estimated using simulated data.

**Figure 9.**
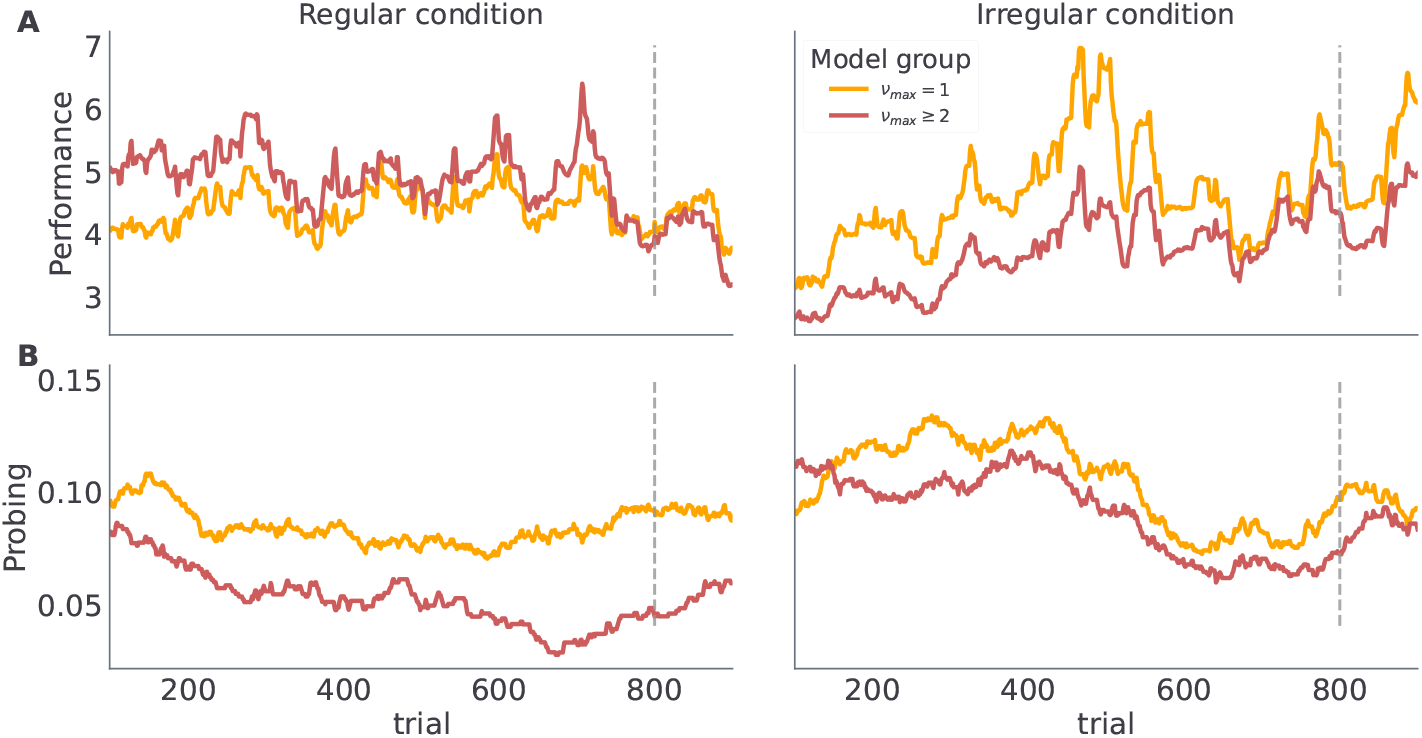
Category based mean estimate of behavioural measures. Each line corresponds to a model class average over behavioural trajectories of subjects assigned to that model class. Note the similarity of the trajectory profiles to the simulated trajectories in ***Figure 5***. **Figure 9–Figure supplement 1.** Subject specific trajectories of behavioural measures.

These findings show a good correspondence between simulated behaviour for differ-ent parametrisation of the model (***v_max_*** = 1 vs ***v_max_*** > 1 in ***Figure 5***), and the participants’ behaviour associated with different model classes (***Figure 9***). There are two possible ex-planations forthis: (i)the model inversion accurately captures the participants behaviour and between-participant sensitivity to temporal regularities of the task, (ii) the group differences come from other free model parameters and do not correspond to differences in sensitivity to temporal structure. To exclude the second option we show in ***Figure 10*** the mean of the posterior estimates of free model parameters *γ*, *p*_−_ and *p*_+_. Note that in both experimental conditions we see a lack of separation between free model parameters associated with each model class.

**Figure 10.**
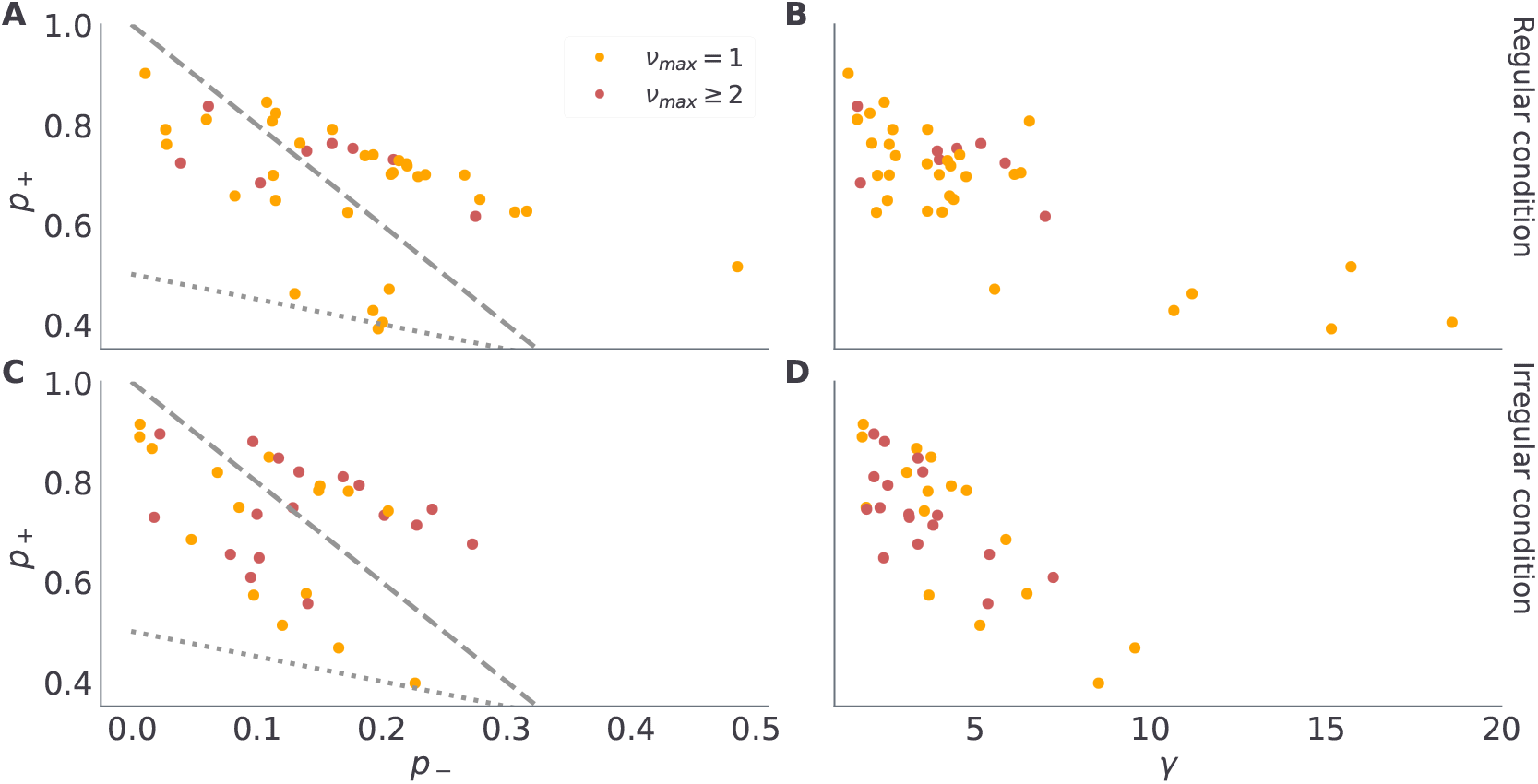
Posterior median of continuous model parameters. Each dot corresponds to the median of the *n* = 1000 samples from the posterior distribution of *γ*, *p*_−_ and *p*_+_ for each participant. The dashed grey lines denotes equality between preferences for losses and epistemic cues, that is, when 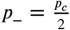 then *p*_−_ = (1 – *p*_+_)/2. In turn, the doted grey line indicates equality between preferences for gains and epistemic cues, that is, when 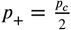 then *p*_−_ = 1 – 3*p*_+_. Note that we use the same colour coding as in the previous figure to denote classification of participants into different model classes.

There are a couple of interesting observations to be made from the posterior expectations of the other model parameters. First, we find in most participants rather large posterior estimates of choice precision *γ*, close to *γ* = 5, suggesting that choice stochasticity is rather low in most participants. Low choice stochasticity means that choices are well aligned with the choice likelihoods encoded in terms of expected free energy (Eq (18)) and the model is rather accurate in predicting behavioural responses. Second, the posterior estimates of outcome preference parameters *p*_−_, and *p*_+_ split subjects in two distinct groups, which correspond to their preference for receiving informative cues when selecting exploratory option. The 29 subjects who never engaged with the exploratory option have a higher preference for losses than for informative cues, hence *p*_−_ ≥ *p_c_*. We marked with the dashed grey line the limiting case of 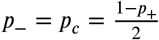, which separates the subjects which did not interact with the exploratory option (above the dashed line) and subjects that were relying on exploratory option to reduce their belief uncertainty (bellow the dashed line). Similarly, the participants who prefer informative cues over gains would have prior preferences over either of the cues in the following region *p_c_* ≥ *p*_+_. The dotted grey line marks the limiting case of 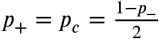.

## Discussion

Sequential activity of neuronal assemblies is one of principled neuronal operations that support higher level cognitive functions ***Eichenbaum (2014)***; ***Buzsáki and Llinás (2017)*** and allow humans to form complex spatio-temporal representation of our every day environment ***Frölich et al. (2021)***. Akin to grid cells known to support representation of both spatial and nonspatial task states ***Fu et al. (2021)***, time cells have been linked to temporal representation of state sequences critical for memory and decision-making ***Eichenbaum (2014)***. Importantly, in spite of these fruitful experimental findings we have no clear computational understanding of how humans learn temporal structure in the service of successfully behavioural adaptation.

Here we introduced a novel computational model of behaviour capable of learning latent temporal structure of a probabilistic reversal learning task with multiple reversals (***Costa et al., 2015***; ***Reiter et al., 2016, 2017***; ***Vilà-Balló et al., 2017***). The computational model combines hidden semi-Markov framework for representing latent temporal structure (***Yu, 2015***) and active inference for resolving exploration-exploitation trade-off (***Friston et al., 2015***, ***2016***). Importantly the model be used for investigating decision making in changing environments in any behavioural task that can be cast as a dynamic multiarmed bandit problem (***Markovic et al., 2021***; ***Gupta et al., 2011***); of which the reversal learning tasks is a special case corresponding to a specific type of two-armed bandit problem.

The probabilistic reversal learning task, which we utilised to demonstrate flexibility of proposed model, is one of the most established paradigms for investigating human behaviour in changing environments and quantifying cognitive disorders. We used model-based analysis of behavioural data to infer temporal expectations of subjects exposed to one of the two task variants: (i) with regular intervals between reversals, (ii) with irregular intervals between reversals. Notably, being able to form expectations about the moment of reversal is critical for achieving high performance in the probabilistic reversal learning task, which we illustrate using simulations. We demonstrated that participants behaviour is highly heterogeneous reflecting the differences in participants expectations about temporal regularities. Importantly, the participants expectations about temporal regularities influence their ability to correctly learn latent temporal structure (especially relevant in the condition with regular between reversal intervals), and is reflected in their performance throughout the experiment.

Importantly, we have extended the standard reversal learning task and incorporated an explicit exploratory option in addition to the two standard options whose choice results in monetary gain or loss. This exploratory option informs the participant about the currently correct choice. The additional behavioural response provides us with more direct access to the individual uncertainty about a correct choice and improves model selection. Interestingly, in addition to participants’ diversity of temporal representation, we find stark differences in their preferences to engage with the exploratory option, suggesting individual differences for the value of information (***Niv and Chan, 2011***) and utilised strategies for resolving the exploration-exploitation trade-off. Critically, their epistemic preferences are not obviously correlated with the quality of the learned temporal structure, as in both groups participants show heterogeneous prior expectations about temporal regularities limiting the available temporal templates, hence the accuracy of temporal representations. However, the willingness to engage with the epistemic options does influence participants’ performance, where higher engagement results in better performance. Therefore, these joint findings reveal distinct components of the computational mechanisms that underlie adaptive behaviour in dynamic environments.

In spite of improved from of the reversal learning, the proposed method still has a limited capability of precisely identifying temporal priors from participants’ responses (see ***Figure 8-Figure Supplement 1***). This introduces uncertainty into the cognitive phenotyping, and constrains how specific we can be about the sources of the participant behavioural variability. We expect that a different experimental paradigm is required to separate participants ability of forming accurate temporal respresentation from the lack of motivation to invest cognitive resources for improved performance. Particularly, in the current reversal learning task it is not possible to separate a range of factors that could contribute to behavioural differences: (i) participants not being able to learn correct latent temporal structure, (ii) participant not expecting that learning higher order statistics can improve performance, and (iii) participants not being motivated enough to invest cognitive resources for performing the task. Being able to decouple contributions of these factors will be critical for applying the behavioural model to cognitive phenotyping and making the approach useful to the research in computational psychiatry and developmental psychology.

To accurately predict future it is critical not only to know that change might be coming but also when the change will occur. To anticipate the changes in our-everyday environments and adapt our behaviour accordingly it is critical accurately estimate and represent elapsed time between relevant events. A range of experimental findings has linked timing of events and hence forecasting the future to underlying Bayesian inference mechanisms (***Jazayeri and Shadlen, 2010***; ***Griffiths and Tenenbaum, 2011***). Most recently ***Maheu et al. (2022)*** has linked sequence learning and prediction in human subjects to an underlying hierarchical Bayesian inference model with distinct hypothesis spaces for statistics and rules corresponding to a set of deterministic temporal templates. The authors conclude that the hierarchical Bayesian inference mechanism underlies human ability to process sequence, similar to hierarchical semi-Markov framework proposed here.

Furthermore, in recent years, various neuroimaging studies have linked different neuro-cognitive domains, such as attention and working memory, to specific spatio-temporal expectations about underlying dynamics of the environment (***Nobre and Van Ede, 2018***). Interestingly, the human ability to estimate and reproduce elapsed time was also previously linked to reward discounting and intertemporal choice behaviour (***Bermudez and Schultz, 2014***; ***Retz Lucci, 2013***; ***Ray and Bossaerts, 2011***). For example, ***McGuire and Kable (2015)*** demonstrated that “impulsivity” (reluctance to wait for a better reward), depends on the hidden statistics of delays — between an initial bad offer and a later but more valuable offer— which human participants experienced. In ***Mikhael and Gershman (2019)*** authors have linked time perception and dopaminergic neuronal activity, demonstrating the role of value-based prediction errors in time representation. Furthermore, time perception and timed behaviour have been linked to all major neuromodulatory systems (***Meck, 1996***) either directly using neuropharmacological manipulations (***Crockett and Fehr, 2014***) or indirectly using neurological disorders (***Story et al., 2016***) and ageing research (***Read and Read, 2004***).

Together these findings provide important evidence for the role of temporal expectations in goal-directed decision making and let one speculate whether a range of aberrant behaviours might be related to an erroneous representation of the temporal structure of the task. Importantly, a computational behavioural models that we introduced here can emulate the learning of temporal structure, hence can become a potent tool linking aberrant behaviour found in cognitive disorders to erroneous prior beliefs about the rules that govern the dynamics of the environment, as suggested by the active inference account of human behaviour (***Friston et al., 2017, 2016, 2015***).

To conclude, the results presented here provide novel insights into computational mechanism underlying the human ability to learn hidden temporal structure of the environment and the computational principles they utilise for making decisions based on temporal representations. The fact that we find behavioural heterogeneity in a population of healthy young adults suggests a potential use of the proposed design and behavioural model for cognitive phenotyping and for revealing causes of aberrant behaviour in clinical populations.

## Methods and Materials

### Ethics statement

All subjects provided written informed consent and were paid on an hourly basis. The Ethical Board of Technical University Dresden approved the study.

### Experiment

#### Probabilistic reversal learning

In the experimental task subjects were deciding between two cards shown on a screen, each showing a different stimulus (a geometric shape, e.g. rectangle, triangle or a question mark) as shown in Figure 1. The reward probabilities associated with the two choice options were anti-correlated on all trials: whenever reward probability of choice A was high (*p_H_* = 0.8) the reward probability of choice B was low (*p_L_* = 0.2), and vice versa. Note that *p_H_* = 1 – *p_L_* on all trials. The location of each stimulus on the screen (right or left side) was kept fixed over trials. After each choice the stimulus was highlighted and depicted for 1.5 s minus the reaction time. The feedback in the form of a gain or a loss was shown for 0.5 s. Similarly, the feedback after an exploratory choice was also shown for 0.5 s. If no response occurred during the decision window of 3s, the message “too slow” was presented, and no outcome was delivered.

All subjects underwent a training session during which they had the opportunity to learn the statistics of the rewards associated with high *p_H_* and low *p_L_* reward probability choices. The set of stimuli used in the training phase differed from the one used during the testing phase. Subjects were instructed that they could either win or lose 10 cents on each trial, and that they will be paid the total amount of money they gained during the testing phase at the end of the experiment. Each subject performed 40 training trials with a single reversal after the 20th trial. Before the start of the testing phase subjects were told that the reward probabilities might change at regular intervals (in both conditions) over the course of the experiment. No other information about reversals or the correlation of choices and outcomes was provided. Thus, the subjects had no explicitly instructed knowledge about the anti-correlated reward probabilities or between-reversal-intervals before the experiment.

Note that, out of *n* = 74 participants *n_p_* = 24 were exposed to the variant of the reversal learning task without epistemic option. This group of subjects belongs to an initial pilot study that used the standard two-choice task design. In the pilot study 14 subjects were assigned to the regular condition and 10 to the irregular condition. We decided to include the subjects from the pilot into the analysis, as we noticed that almost 30% of subjects, in the post pilot group, choose not to interact at all with the exploratory option, even when that was a possibility. We will not explore this finding here in more detail, but we can exclude their misunderstanding of the task as a potential confound, as we provided a detailed instructions and training before they performed the task (see Experiment for more details).

#### Behavioural measures

To quantify behaviour we have used two summary measures: (i) performance, defined as odds of making a correct choice, and (ii) probing, defined as odds of making an exploratory choice.

The process of computing *performance* is illustrated in ***Figure 11***. We first label subjects’ responses as either correct or incorrect, depending on whether a card with higher reward probability was selected or not, see ***Figure 11** A*. Then we compute a probability of making a correct choice within a 201 trial window, centred at the current trial number *t*, see ***Figure 11**B*. Finally, for each trial we compute performance as odds of being correct, see ***Figure 11**C*.

**Figure 11.**
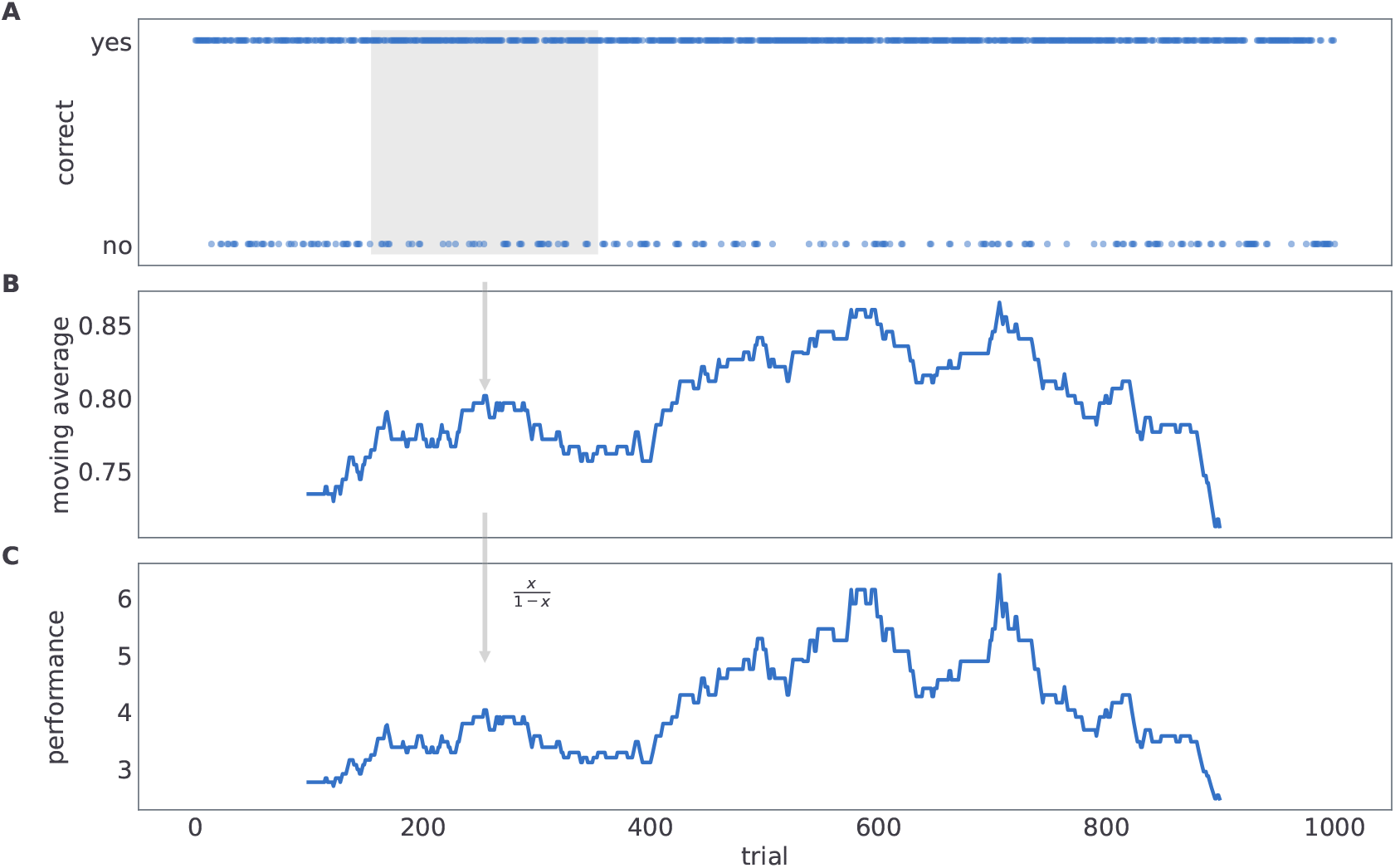
Computing behavioural performance. The process of computing *performance*. **A**. We label subjects’ responses as either correct or incorrect, depending on whether a card with higher reward probability was selected or not. **B** We compute a probability of making a correct choice within a 200 trial window, centred at the current trial number *t*. **C** For each trial we compute performance as odds of being correct.

The *probing* is computed in similar manner to performance, with the only difference that we label choices as either exploratory or exploitative depending on whether subjects have chosen the exploratory option (middle card in ***Figure 1***), or not. Probing is defined as the odds of selecting the exploratory option within a 200 trials time window.

#### Behavioural model

To introduce the generative model of task dynamics, and subsequently derive the behavioural model via model inversion methods, we will consider the following features of the task. At any trial the task environment is in one of the two possible states, defined as the configuration of reward contingencies. For example, state one corresponds to stimulus A being associated with a high reward probability *p_H_*, and state two to stimulus B being associated with a low reward probability *p_L_*. Subjects do not know in advance how likely rewards and losses are when making a correct choice compared to making an incorrect choice, and this is something they have to learn during the course of experiment. In other words, we also treat reward probabilities (*p_H_*, and *p_L_*) as latent variables. Between trials the state can change, i.e. when a reversal occurs but only after a certain minimum number of trials has elapsed since the last state change. Depending on the experimental condition the between reversal duration will either be semi-regular (occurring every 20 trials with small variability) or irregular (occurring every 20 trials, but with maximal variability)

The explicit representation of state duration *d* enables us to associate changes in state transition probabilities with the current trial and the moment of the last change. The dependence of state transition probability on the number of trials since the last change corresponds to the formalism of hidden semi-Markov models (HSMM) (***Yu, 2010***; ***Murphy, 2002***), which allows mapping complex dynamics of non-stationary time series to a hierarchical, time aware, hidden Markov model. However, using an explicit representation of context duration is inefficient, as it requires an enormous state space representation. Here, we will instead adopt a phase-type representation of duration distribution (***Varmazyar et al., 2019***) which substitutes duration variable *d* ∈ {1,…, ∞} with a phase variable *f* ∈ {1,…, *f_max_*}, allowing for a finite state representation of an infinite duration state space.

In what follows we will define the components of the generative model (observation likelihood, the dynamics of latent variables, and the parametrisation of the dynamics) and derive the corresponding update rules for latent variables and state, hence enabling the learning of different temporal contexts during the experiment. The graphical representation of the generative model is shown in ***Figure 4***.

Practically we introduce four latent states, to describe the task on any trial:

- First, the configuration of reward contingencies can be in one of the two possible states. Hence, 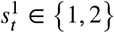 which describes which card is associated with high reward probability and which with low reward probability.
- Second, choosing one of the options on a given trial corresponds to setting the task in one of the three possible choice states 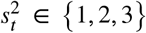 (chosen left card, chosen middle card - exploratory option, and chosen right card) corresponding to the chosen option. The choice of the option is deterministic and this state is always known with certainty after the choice is made.
- Third, current phase *f_t_* ∈ {1,…, *v* + 1} of the task dynamics. The phase latent variable controls transitions of latent state 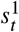, where the change of state is only possible if the end phase (*f_t_* = *v* + 1) is active on the current trial. Note that the larger the number of phases is (parameter *v* ∈ 1,…) the more regular is the occurrence of reversals. We have limited here the number of phases by setting *v* = 10, as this is sufficiently large for accurate representation of reversal dynamics in regular condition.
- Fourth, temporal template *m*. Latent temporal template defines the frequency of reversals, *μ* (mean between-reversal duration) and the number of latent phases *v*, that is the regularity of reversals.

##### Observation likelihood

The observation likelihood links latent states 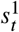, and 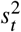) with probabilities of observing different possible outcomes in those states.

In the temporal reversal learning task there are four possible outcomes: (1) loss of 10 Eurocents, (2) gain of 10 Eurocents, (3) the correct card is left card, or (4) the correct card is the right card. Therefore, we define the observation likelihood as a categorical distribution

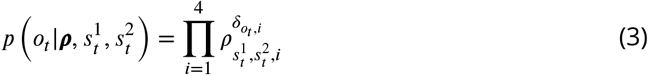

where *i* denotes the outcome type, *o_t_* ∈ {1,…, 4}. The probabilities of different outcomes are parameterised via 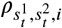, where each state tuple 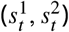 corresponds to a unique probability of observing any of four possible outcomes. We define prior beliefs about outcome probabilities in the form of a product of Dirichlet distributions

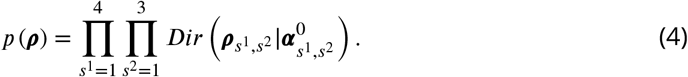

We set the parameters of Dirichlet priors to the following values:

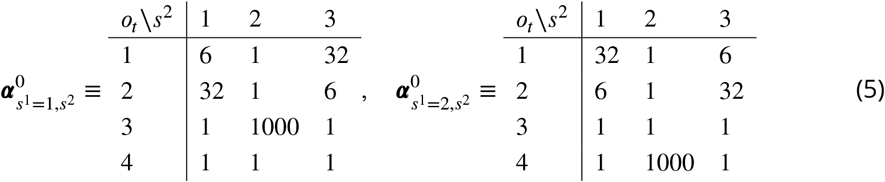

The above configuration for the parametrisation of prior Dirichlet probabilities reflects an assumption that the participants have formed during training an initial - vague beliefs - about reward probabilities associated with different actions in different states. We assume that participants are highly certain that selecting the epistemic option does not return gain or loss (high value of ***α***_*s*^1^, *s*^2^=2_ for the corresponding outcome in both states). Furthermore, we assume that participants have formed good expectations gain/loss probabilities (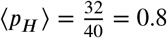, and 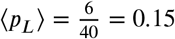), but that they are still uncertain about the exact values. Weak priors about outcome probabilities allow for ongoing adaptation of beliefs during the course of experiment.

##### Hidden state dynamics

To formalise the presence of sequential reversals, we define the phase dependent state transition probability as follows

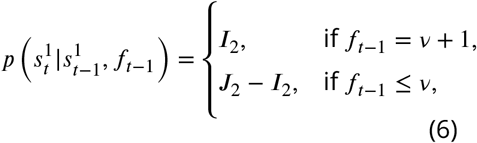

where *I*_2_ denotes the 2 × 2 identity matrix and *J*_2_ denotes the 2×2 all-ones matrix. The above relations describe a simple deterministic process for which the current state 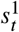 remains unchanged as long as the phase variable *f*_*t*−1_ remains below the end phase, *v* + 1. The transition between states occurs with certainty (e.g. if 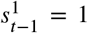 then 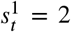) once the end phase is reached, that is, when *f*_*t*–1_ = *v* + 1.

Although it is possible to condition state changes on a duration variable *d*, as demonstrated in ***Marković et al. (2019)***, such an explicit representation is inefficient as it requires large state spaces (***Yu and Kobayashi, 2003***; ***Vaseghi, 1995***). Here we adopt the discrete phase-type (DPH) representation of duration distribution (***Varmazyar et al., 2019***). The DPH representation defines transitions between phase variables *f_t_* and the following parametrisation of phase transition probabilities corresponds to the DPH representation of the negative binomial distribution

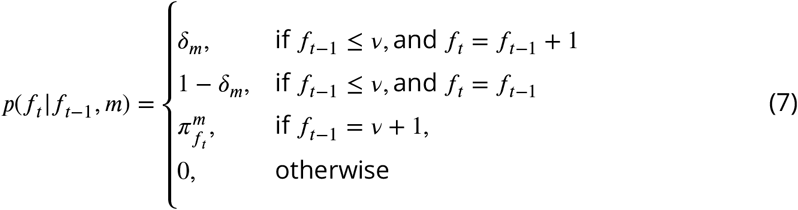

where 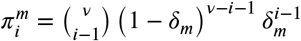 for *i* < *v* + 1 and 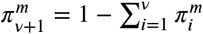.

The corresponding negative binomial distribution of between-reversal duration can be expressed as follows

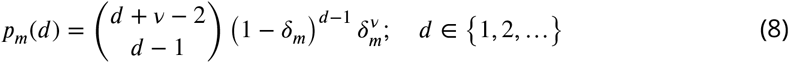

where the expected duration corresponds to

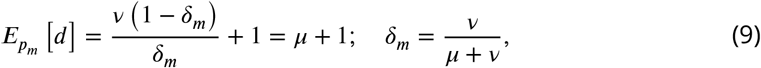

and variance, hence uncertainty about duration regularity, to

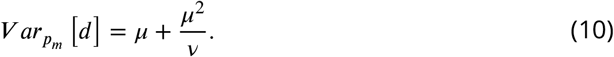

Note that the parameter *v* of the negative binomial distribution, acts as a precision parameter. We illustrate this in Figure 12.

The choice of prior beliefs about the between-reversal interval *d* in the form of a negative binomial distribution has interesting consequences on the dynamics of the marginal probability that a reversal will occur at some future point *τ*

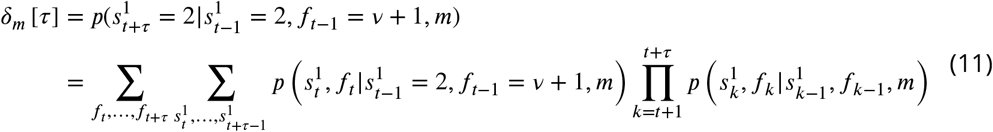

In Figure 13 we show the dependence of the future reversal probability *δ_m_* [*τ*] on the precision parameter *v*, given a fixed mean duration *E_p_m__* [*d*] = 20. Note that for *v* = 1 we get a constant transition probability, which corresponds to the expectations of change probabilities found in hidden Markov models. In contrast, for larger values of *v* one obtains a trial-dependent, effective transition probability with values alternating between low and high probabilities in a periodic manner. This temporal dependence of the transition probability will affect the inference process. The agent will become insensitive to subsequent reversals occurring a few trials after the previous reversal, and highly sensitive to reversals occurring twenty to thirty trials after the previous reversal.

**Figure 12.**
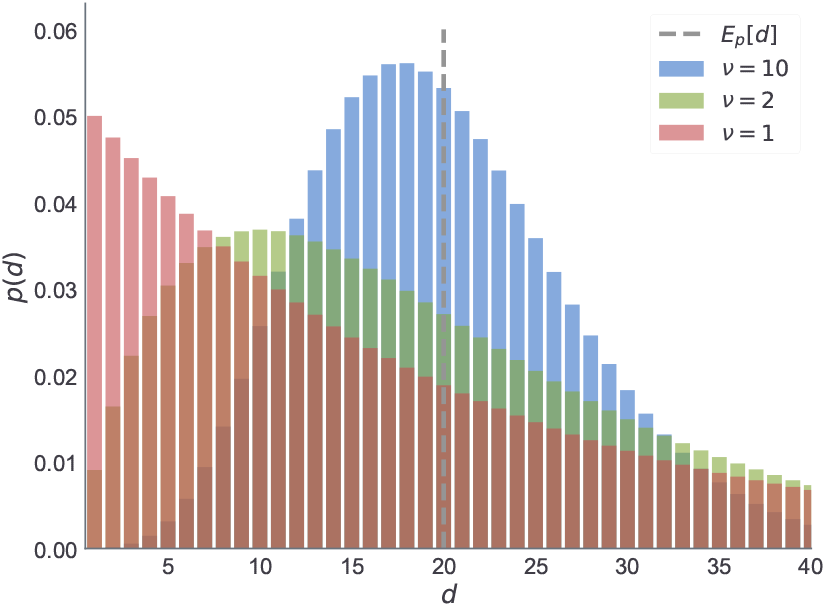
Negative binomial distribution. We illustrate here the changes in the negative binomial distribution as a function of shape parameter *v* which is inversely proportional to the variance of between reversal durations. Note that for higher values of *v* the distribution peaks around its expected value (dashed line). As the variance increases (green) the mode shifts toward zero. The limiting case of the negative binomial distribution in the form of geometric distribution (red) corresponds to *v* = 1. For all three cases we fixed the mean duration to the same value.

**Figure 13.**
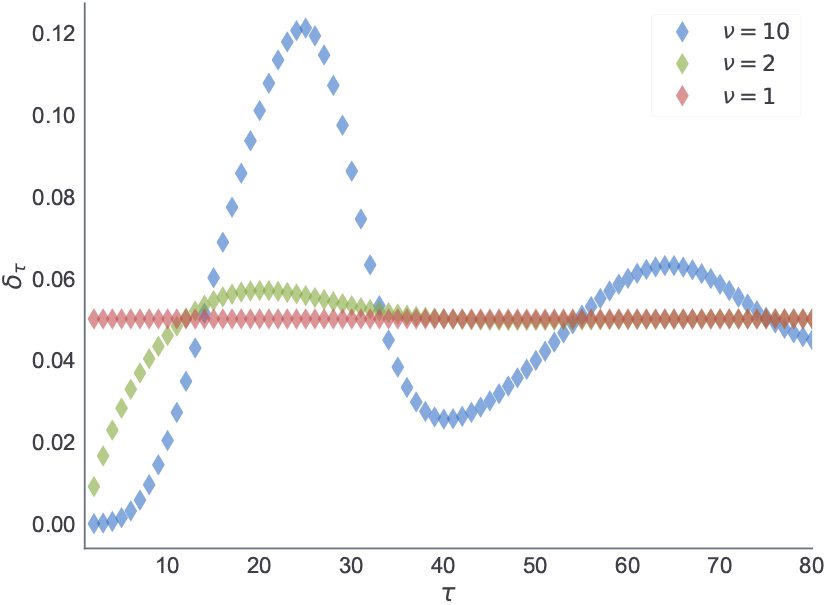
Expected transition probability at future trial *τ*. Estimate of the transition probability *δ_m_*[*τ*], Eq (11), at a future trial *τ* conditioned upon a reversal at *t* and known initial state 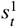. Each curve corresponds to estimates of the transition probability obtained from prior beliefs *p_m_*(*d*) shown in Figure 12.

Finally, the choice states 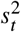 are fully dependent on the current choice *a_t_* ∈ {1, 2, 3}, and we express the state transition probability as

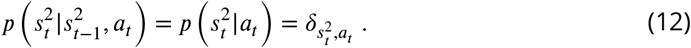

In practice this means that the agent is always certain about the choice it made and how that choice impacted the state of the task. Therefore, the posterior estimate over 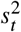 can be trivially expressed as

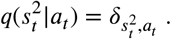

##### Active Inference

In active inference, agents form posterior beliefs both about latent states of the environment and about their own actions. In other words, both perception and action selection are cast as inference problems (***Botvinick and Toussaint, 2012***; ***Attias, 2003***). Practically, we will use variational inference for defining update rules for beliefs (***Friston et al., 2017***; ***Blei et al., 2017***). In what follows we will first introduce perception as minimisation of the variational free energy (upper bound on log-marginal likleihood) with respect to posterior beliefs over latent states, and after that introduce action selection as minimisation of the expected free energy (***Smith et al., 2022***), that is, expected surprisal about future outcomes.

We write the generative model of outcomes *o_t_* on trial *t* as

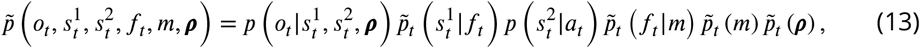

where we use 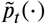 to denote prior beliefs conditioned on a sequence of past outcomes, *o*_1:*t*–1_ = (*o*_1_,…,*o*_*t*–1_) and choices *a*_1:*t*–1_ = (*a*_1_,…, *a*_*t*–1_. Given a choice 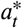 and an observed outcome *o_t_* at trial *t*, the approximate posterior belief *q_t_*(*x*) over latent states 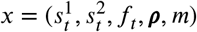 is obtained in two steps:

- We first compute the marginal likelihood with respect to 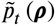, and obtain the exact marginal posterior over discrete states using the Bayes rule

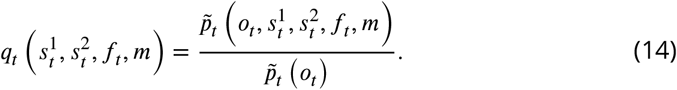
- Given the marginal posterior 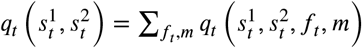 we compute the posterior over outcome probabilities using the variational message passing update

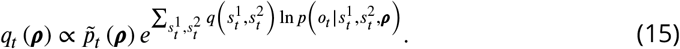

As we initially defined the prior over outcome probabilities in the form of a Dirichlet distribution with parameters ***α*^0^**, we can express the posterior estimate on every trial in the same functional form. Hence,

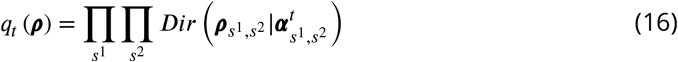

where

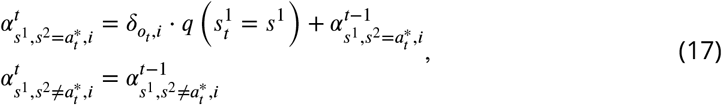

and 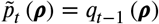. The above belief updating scheme corresponds to the variational surprise minimization learning algorithm (***Liakoni et al., 2021***; ***Markovic et al., 2021***) adapted to the categorical likelihood and the Dirichlet prior.

##### Action selection

In active inference, decision strategies (behavioural policies) are chosen based on a single optimisation principle: minimising expected surprisal about observed and future outcomes, that is, the expected free energy (***Smith et al., 2022***; ***Schwartenbeck et al., 2019***).

Here, we will express the expected free energy of a choice *a* on trial *t* as

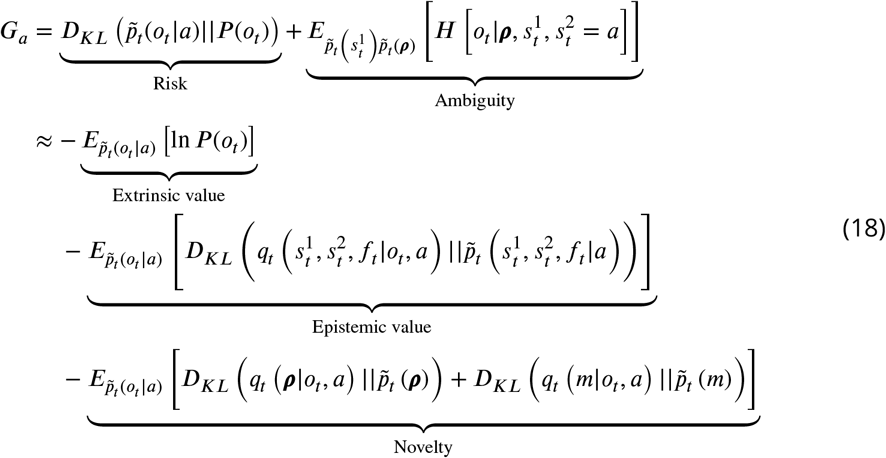

where *P*(*o_t_*) denotes prior preferences over outcomes, 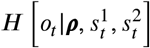 the entropy of out-come likelihood 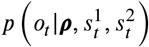, and *D_KL_*(*p*||*q*), stands for the Kullback-Leibler divergence between two probability densities: *p* and *q*. Note that action selection based on minimisation of expected free energy would have an implicit dual imperative (see the different factorisations in Eq (18)): On one hand, the expected free energy combines ambiguity and risk. On the other hand, it consists of information gain (epistemic value + novelty) and extrinsic value. Therefore, selecting actions that minimise the expected free energy dissolves the exploration-exploitation trade-off, as every action contains both expected value and information gain. This is a critical feature of action selection which allows us to account for epistemic choices as used in our experimental paradigm (see Figure 1).

At any trial *t* choice *a_t_* is sampled from choice beliefs *p*(*a_t_*) (cf. planning as inference ***Botvinick and Toussaint*** (***2012***); ***Attias*** (***2003***)) defined as

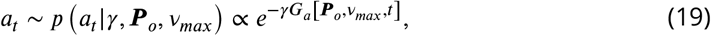

where parameter *γ* corresponds to choice precision, which we will attribute to empirical choice behaviour of participants. Therefore, for describing participants’ behaviour we assume that the action selection process is corrupted by external sources of noise;e.g. mental processes irrelevant for the task at hand. In our simulations we will fix *γ* to a reasonably large value, to achieve approximate free energy minimisation as the following relation will be satisfied

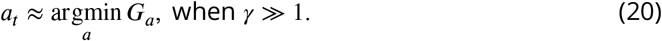

Notably, here we consider the simplest form of active inference in which expected free energy is computed from a one-step-ahead prediction. This is a standard simplification for environments in which actions cannot interfere with the state transitions, as is the case in typical dynamic multi-armed bandit problems (***Markovic et al., 2021***).

To express the expected free energy, *G*(*a_t_*), in terms of beliefs about arm-specific reward probabilities, we will first constrain the prior preference to the following categorical distribution

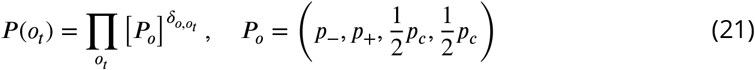

In active inference, prior preferences determine whether a particular outcome is attractive, that is, rewarding. Here we assume that all agents prefer gains (*o_t_* = 2) over losses (*o_t_* = 1). Hence, we constrain parameter values such that *p*_+_ > *p*_−_ holds always. The ratio 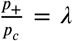 determines the balance between epistemic and pragmatic imperatives. When prior preferences for gains are very precise, corresponding to large *λ*, the agent becomes risk sensitive and will tend to forgo exploration if the risk is high;see Eq. 18. Conversely, a low lambda corresponds to an agent which is less sensitive to risk and will engage in exploratory, epistemic behaviour, until it has familiarised itself with the environment.

Given the following expressions for the marginal predictive likelihood,

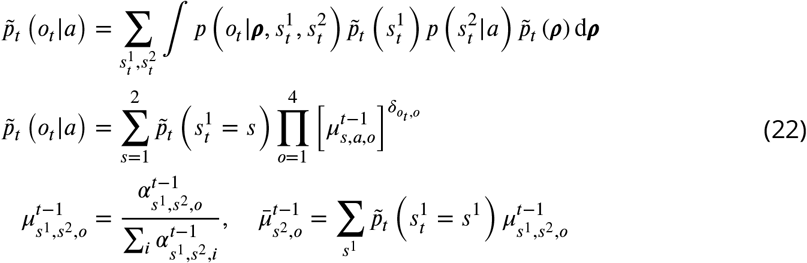

we get the following expressions for the expected free energy

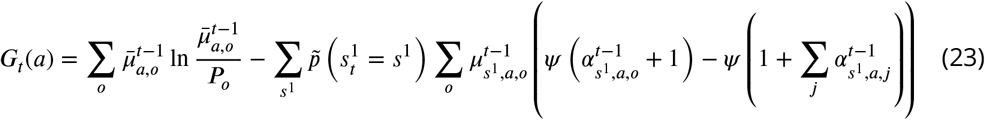

Above we have used the following relation

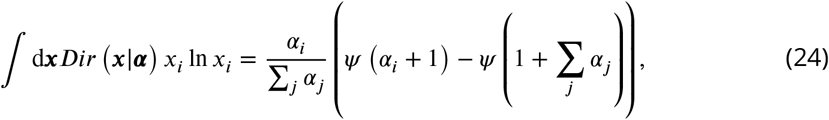

for computing ambiguity term in Eq (18).

#### Model inversion

Given a sequence of subjects’ responses 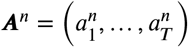, where *n* denotes subject index and *T* = 1000 denotes the total number of trials, the response likelihood is defined as

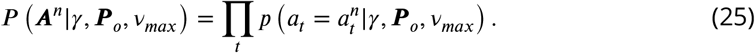

Note that for estimating the posterior over model parameters (*γ*, ***P**_o_*, *v_max_*) we ignore the first 400 responses from the likelihood. We expect that during these first trials, subjects are still getting used to the task, and potentially use additional strategies for representing the task and making choices. As we do not model all possible task representations, exclusion of initial trails reduces the noise in model comparison. Importantly, we do use the entire set of responses for computing belief trajectories of the active inference agents, that is, we expose the agent to the complete sequence of individual responses and the corresponding outcomes.

For the posterior estimates from the behavioural data we used the following parametric generative model of behaviour, a mixture model:

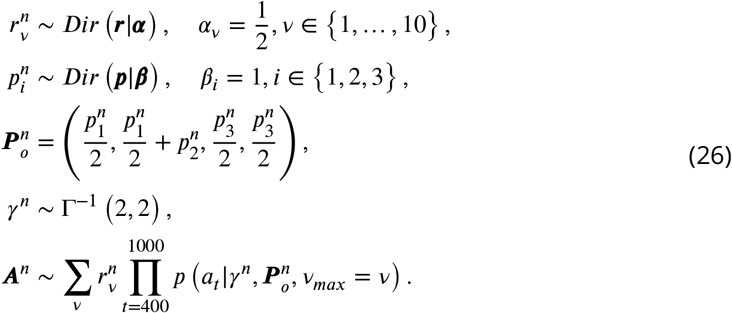

This mixture model unifies parameter estimation with model comparison, as marginal posterior estimates 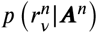 corresponds to the posterior probability of the precision parameter *v_max_*. To classify subjects’ behaviour in terms of adaptability of temporal representations we use the exceedance probability of the marginal posterior defined as

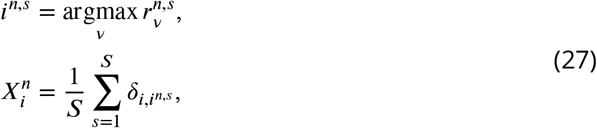

where 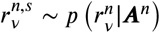 denotes a sample from the posterior. Finally, the most likely precision parameter 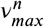 of the *n*th subject corresponds to

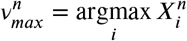

We used the probabilistic programming library Numpyro (***Phan et al., 2019***) for implementing the full generative model. Numpyro library provides an interface to multiple state-of-the-art inference schemes. For drawing samples from the posterior we have used Numpyro’s implementation of the No-U-Turn sampler (NUTS) (***Hoffman et al., 2014***). NUTS is an self-tuning version of the Hamiltonian Monte Carlo, a popular Markov Chain Monte Carlo algorithm for avoiding random walks and sensitivity to between-parameter correlations.

## Acknowledgments

The authors thank Karl Friston for valuable suggestions and discussions.

## Funding

The study was supported by by the German Research Foundation (DFG, Deutsche Forschungsgemeinschaft), SFB 940/3 - Project A09 awarded to SS and SFB 940/3 - Project B7 awarded to AMFR. SS acknowledges further support by DFG TRR 265/1 (Project ID 402170461, B09) and Germany’s Excellence Strategy - EXC 2050/1 (Project ID 390696704) - Cluster of Excellence “Centre for Tactile Internet with Human-in-the-Loop” (CeTI) of Technische Universität Dresden. AMFR acknowledges further support by the German Research Foundation (DFG RE 4449/1-1, RTG 2660-B2) and by a 2020 BBRF NAR-SAD Young Investigator Grant from the Brain and Behavior Research Foundation. This study was partially funded by funding opportunities for young scientists (Anschubfinanzierung) from the Department of Psychology of the Technische Universität Dresden.

---

**Figure 7-Figure supplement 1.**
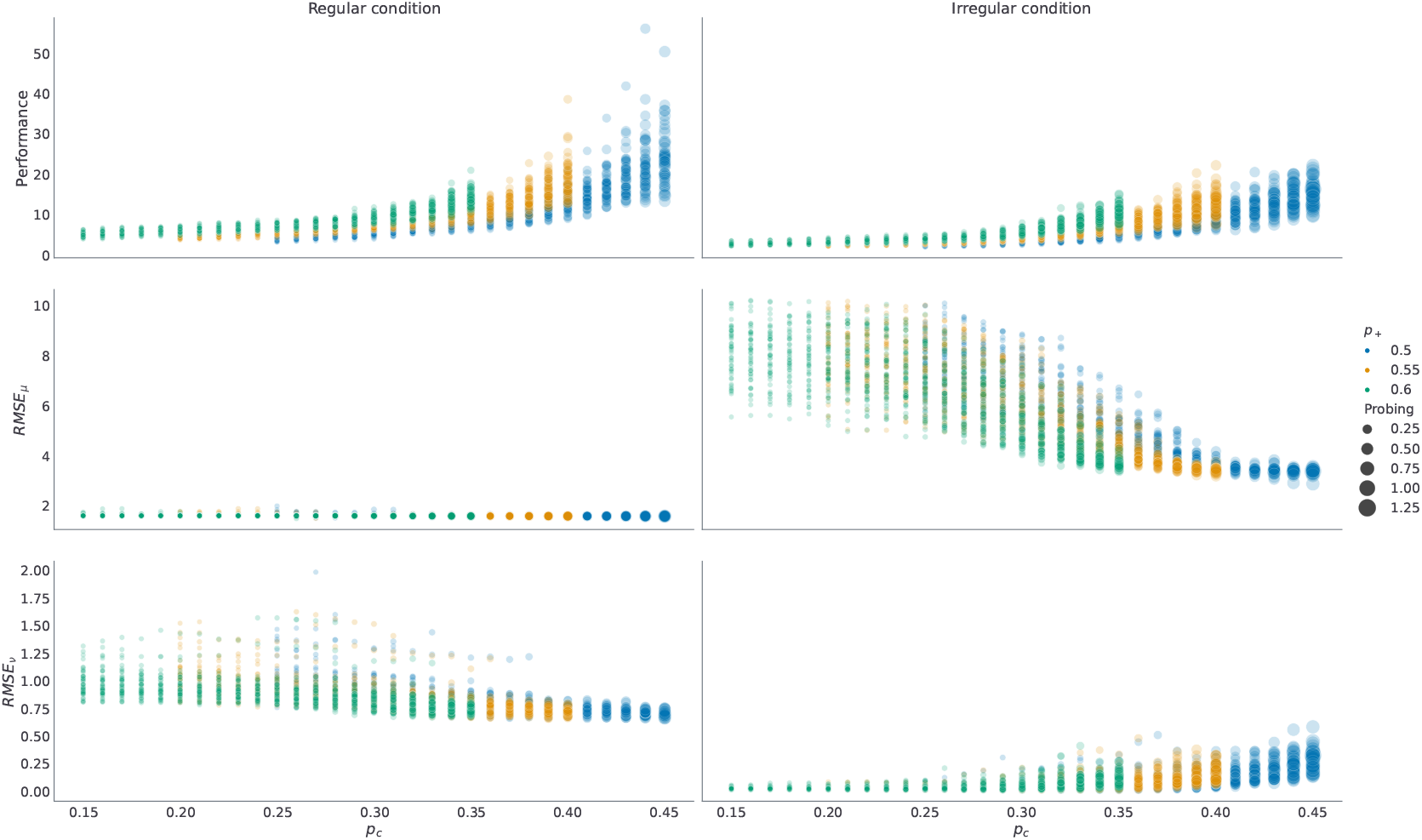
As in the main figure each line corresponds to an average over *n* = 50 simulated trajectories with *v_max_* = 10 and *γ* = 5.

**Figure 8-Figure supplement 1.**
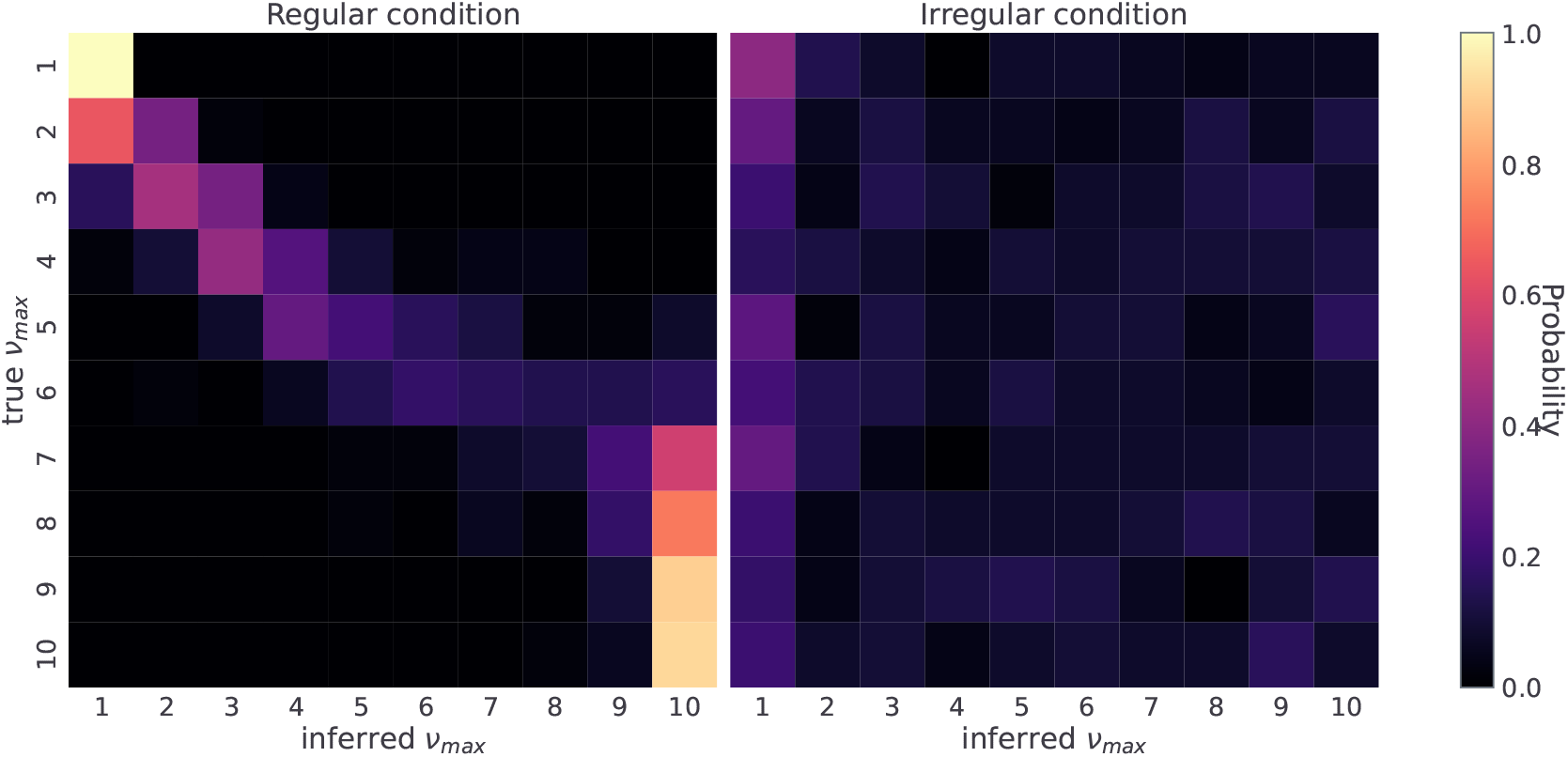
Probability of assigning simulated behavioural responses to possible *v_max_* values given the true *v_max_* ∈ {1,…, 10}. The confusion matrix was estimated based on *n* = 100 simulated responses (50 simulated subjects in each condition) for each generative *v_max_* value. In all simulation the remaining free model parameters were fixed to *γ* = 5, and ***P**_o_* = (0.1, 0.6, 0.15, 0.15).

**Figure 9-Figure supplement 1.**
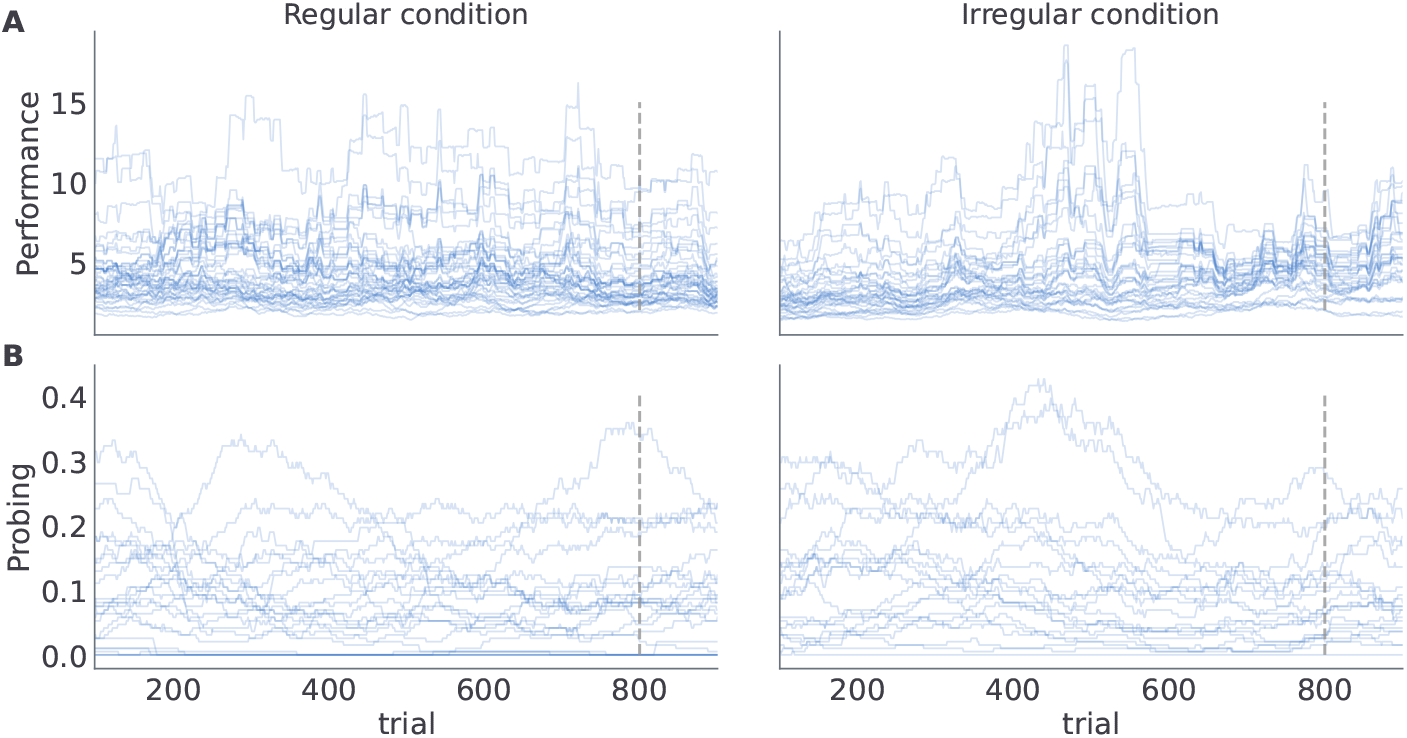
Each line corresponds to a trajectory of: **A** performance, and **B** probing for individual subjects.

## Notes

https://github.com/dimarkov/pybefit/tree/master/examples/temporal_rev_learn

### Competing Interest Statement

The authors have declared no competing interest.

